# Sigh breathing rhythm depends on intracellular calcium oscillations in a population of inspiratory rhythmogenic preBötzinger complex neurons in mice

**DOI:** 10.1101/2022.05.05.490664

**Authors:** Daniel S. Borrus, Cameron J. Grover, Gregory D. Conradi Smith, Christopher A. Del Negro

## Abstract

The preBötzinger Complex (preBötC) of the lower brainstem generates two breathing-related rhythms: one for inspiration on a timescale of seconds and another that produces larger amplitude sighs on the order of minutes. Their underlying mechanisms and cellular origins remain incompletely understood. We resolve these problems via a joint experiment and modeling approach. Blocking purinergic gliotransmission does not perturb either rhythm and imaging experiments show that both rhythms emanate from the same glutamatergic neuron population. We hypothesized that these two disparate rhythms emerge in tandem wherein recurrent excitation gives rise to inspiratory rhythm while a calcium oscillator generates sighs; there is no obligatory role for gliotransmission, hyperpolarization activated mixed cationic current (*I*_h_) in neurons, or synaptic inhibition-mediated coupling of separate populations. We developed a mathematical model that instantiates our working hypothesis. Tests of model predictions validate the single-population rhythmogenic framework, reproducing disparate breathing-related frequencies and the ability for inspiratory and sigh rhythms to be separately regulated in support of respiration under a wide array of conditions. Here we show how a single neuron population exploits two cellular tool-kits: one involving voltage-dependent membrane properties and synaptic excitation for inspiratory breathing (eupnea) and an intracellular biochemical oscillator for sighs, which ventilate and maintain optimal function in the compliant mammalian lung.

**SIGNIFICANCE STATEMENT:** Breathing consists of two vital rhythms: one for eupnea that serves periodic physiological gas exchange and the other for sighs, which are larger breaths that occur minutes apart and serve to optimize pulmonary function. These rhythms with disparate frequencies emerge via a mechanism that is simpler than previously envisaged: it results from one neuron population (not two as previously thought) without need for gliotransmission or synaptic inhibition-mediated coupling of neuronal populations. We show that a low-frequency intracellular calcium oscillation underlies sighs and functions in parallel with the higher-frequency voltage-dependent network oscillation that drives eupnea. Exploiting two separate cellular tool kits enables quasi-independent breathing rhythms, which are unique features of breathing in mammals with compliant lungs.

## Introduction

Central pattern generator (CPG) circuits produce the underlying rhythm and rudimentary motor pattern for rhythmic behaviors (1). In mammals, depending on context, the locomotor CPG produces either walking, running, or bounding where each form of locomotion is mutually exclusive (2); the oromotor CPG produces either chewing, lapping, or swallowing where, again, one form of ingestive behavior precludes the others (3, 4). The breathing CPG, however, generates two rhythms in tandem: one for eupnea to ventilate the lungs on a second-to-second basis (∼3 Hz in rodents to ∼0.2 Hz in humans) and another for sighs to optimize pulmonary function with periodicity on the order of minutes (0.5-5 min^-1^ in rodents to ∼0.2 min^-1^ in humans) (5). We investigated their underlying mechanisms to elucidate the neural origins of breathing and advance understanding of CPGs in general.

Eupnea and sigh are both forms of inhalation and both emanate from the preBötzinger Complex (preBötC) of the lower brainstem (6, 7). The active phase of eupnea, inspiration, initiates due to recurrent excitation among glutamatergic preBötC interneurons (8, 9) and terminates due to refractory mechanisms including synaptic depression and recruitment of neural activity-dependent outward currents (10–12).

The sigh mechanism is less clear but may involve calcium (Ca^2+^) oscillations because it is relatively voltage-insensitive in *in vitro* models of breathing and can be disrupted by antagonists of voltage-gated Ca^2+^ channels, intracellular Ca^2+^ release, or Ca^2+^ chelation (7, 13, 14). One cogent model of the embryonic system posited a discrete neuron population that generates sigh rhythm via Ca^2+^ oscillations that depend on hyperpolarization-activated mixed cationic current (*I*_h_). Also, a discrete neuron population generating inspiratory rhythm via voltage-dependent persistent Na^+^ current (*I*_NaP_) putatively synchronizes with the sigh oscillator through chloride-mediated synaptic inhibition (13). The veracity of the model depends on: i) the existence of dedicated sigh and inspiratory neuron populations, ii) *I*_h_ being sigh rhythmogenic, iii) *I*_NaP_ being inspiratory rhythmogenic, and iv) synaptic inhibition coupling the two populations. Yet, all four stipulations are problematic.

First, the literature is contradictory regarding whether dichotomous eupnea- and sigh-specialized neuron populations exist in the preBötC (15, 16); furthermore, a more recent report suggests that preBötC astrocytes are sigh rhythmogenic (17). Second, experimental data are unsupportive regarding a rhythmogenic role for *I*_NaP_ in inspiration (18–22). Third, blocking *I*_h_ stops sigh rhythm in the embryonic preBötC (13) but this observation has not been corroborated in the preBötC postnatally. Lastly, synaptic inhibition is not necessary to coordinate inspiratory and sigh oscillations in the slice model of rhythmogenesis postnatally (23).

We exploited rhythmically active slice preparations postnatally to address the unresolved issues recapped above, constructed a mathematical model of inspiratory and sigh rhythmogenesis, and then evaluated its testable predictions. We posit that a recurrent excitation-based network oscillator generates inspiration while interactions between plasma membrane (PM) Ca^2+^ fluxes and Ca^2+^ excitability of endoplasmic reticulum (ER) drive sighs. Disparate mechanisms enable inspiratory and sigh rhythms to operate quasi-independently within a single neuronal population tasked with maintaining functionality of compliant mammalian lungs and adjusting breathing for different physiological contexts.

## Results

### Inspiratory and sigh rhythms can be separately modulated

Slice preparations that retain the preBötC remain rhythmically active *in vitro* and generate breathing related motor output via the hypoglossal (XII) nerve. We monitored inspiratory and sigh frequency via preBötC field recordings and XII motor output while manipulating the baseline membrane potential of preBötC neurons via artificial cerebrospinal fluid (aCSF) extracellular K^+^ concentration ([K^+^]_o_) (Fig. 1A). Mean inspiratory frequency measured 0.14 ± 0.05 Hz (N = 19) at 9 mM [K^+^]_o_ aCSF. Decreasing [K^+^]_o_ slowed inspiratory frequency incrementally such that at 3 mM [K^+^]_o_ it measured 0.009 ± 0.011 Hz (N = 9) or zero (N = 11) (Fig. 1B, left). These data align with previous studies showing that baseline membrane excitability governs inspiratory frequency (24, 25), consistent with a recurrent excitation-based network oscillator as its core underlying mechanism (26–28, 12). Mean sigh frequency measured 0.66 ± 0.17 min^-1^ (N = 13) at 9 mM [K^+^]_o_ (Fig. 1B, right). Decreasing [K^+^]_o_ slowed the sigh frequency incrementally such that at 3 mM it measured 0.26 ± 0.08 min^-1^ (N = 5). Sigh rhythm was 19-fold less sensitive to changes in [K^+^]_o_ than inspiratory rhythm (relative frequency increase of m = 19, r^2^ = 0.94, p = 3.6 × 10^−4^) (Fig. 1C).

**Figure 1:**
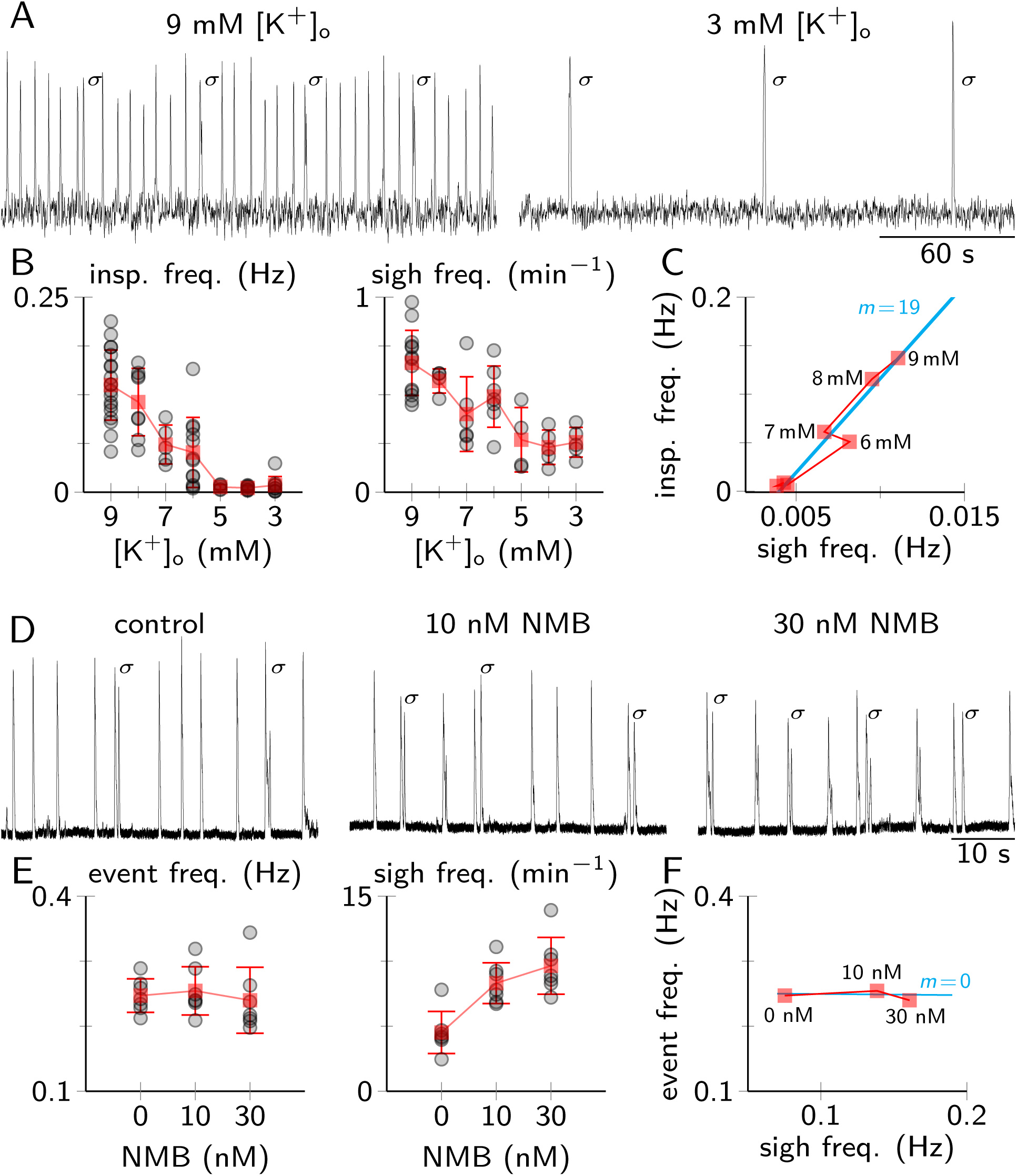
Differential modulation of inspiratory and sigh rhythm frequency. (A) preBötC field recording capturing inspiratory and sigh rhythms at 9 and 3 mM extracellular K^+^ concentration in the aCSF ([K^+^]_o_). Sigh events are indicated by σ. (B) Inspiratory and sigh frequencies plotted as a function of aCSF [K^+^]_o_. Grey circles show individual slices; red squares show mean frequency (N = 19 inspiratory, N = 13 sigh). (C) Mean inspiratory frequency plotted versus corresponding mean sigh frequency for different aCSF [K^+^]_o_. (D) XII nerve recordings in control and after Neuromedin-B (NMB) application. (E) Event (inspiratory) and sigh frequencies plotted at different NMB doses. Grey circles represent individual slices; red squares show mean frequency (N = 8). (F) Event (inspiratory) and sigh frequencies plotted at different NMB doses.

Neuromedin B (NMB) is a bombesin-like peptide associated with sigh regulation (29) (Fig. 1D). We meta-analyzed NMB effects on inspiratory and sigh frequency from Li *et al*. (28). Sigh frequency increased from 4.5 ± 1.6 min^-1^ in control to 8.3 ± 1.6 min^-1^ following 10 nM NMB and to 9.7 ± 2.2 min^-1^ following 30 nM NMB (r^2^ = 0.58, p = 5.8 × 10^−5^) (Fig. 1E, right). We analyzed inspiratory frequency via all *events*, i.e., both standalone inspiratory bursts and sigh bursts, which typically build off an inspiratory burst with very short latency (≲ 1 *s*) (7, 23). Control inspiratory (event) frequency of 0.25 ± 0.03 Hz remained unaffected by 10 nM NMB (0.25 ± 0.04 Hz) and 30 nM NMB (0.24 ± 0.05 Hz) (Fig. 1E, left). Comparing the change in inspiratory (event) frequency to the change in sigh frequency yields a flat line (m = 0, r^2^ = 6.4 × 10^−3^, p = 0.73) (Fig. 1F), indicating that sigh rhythm can be modulated without changing inspiratory rhythm frequency (14). These data suggest that inspiratory and sigh rhythms have different mechanisms.

### Purinergic signaling is not necessary for sigh rhythm generation

One possibility is that two discrete rhythmogenic mechanisms are embodied in different cell populations, neural and/or non-neural (17). Astrocytes can generate intracellular Ca^2+^ oscillations and gliotransmission via purinergic P2 receptors modulates inspiratory preBötC rhythms (30–32), which suggests astrocytes communicating with preBötC neurons via purinergic P2 receptors is a feasible mechanism for sigh rhythmogenesis.

We monitored preBötC neurons derived from progenitors that express the transcription factor *Dbx1* (hereafter: Dbx1 neurons), which comprise the inspiratory rhythmogenic preBötC core (33–35). Multiphoton imaging of membrane-bound GCaMP6f in Dbx1;Ai148 mouse slices produced measurable Ca^2+^ transients simultaneously recorded with XII motor output (Fig. S1A,B). Blocking the spectrum of P2 receptors via a cocktail of antagonists (50 μM PPADS, 50 μM suramin, 10 μM TNP-ATP, 10 μM MRS2179, and 10 μM MRS2578) did not modify the frequency of either inspiratory rhythm (0.27 ± 0.06 Hz in control vs. 0.26 ± 0.07 Hz in P2 antagonist cocktail, p = 0.41) or sigh rhythm (0.73 ± 0.22 min^-1^ in control vs. 0.74 ± 0.18 min^-1^ in P2 antagonist cocktail, p = 0.87) (Fig. S1C). A parallel experiment employed the highly selective P2Y_1_ antagonist MRS2279 because recent evidence in preprint form purports that gliotransmission specifically via P2Y_1_ receptors is obligatory for sigh rhythmogenesis *in vitro* (17). Bath-applied 20 μM MRS2279 had no effect on either inspiratory or sigh rhythm (inspiratory rhythm: 0.20 ± 0.04 Hz in control vs. 0.26 ± 0.06 Hz in MRS2279, p = 0.13; sigh rhythm: 0.86 ± 0.45 min^-1^ in control vs. 0.61 ± 0.28 min^-1^ in MRS2279, p = 0.26) (Fig. S2A-C).

We also plotted a histogram of inter-event intervals (Figs. S1D and S2D). In control there are peaks at 5 s and ∼10 s that reflect the typical interval between inspiratory bursts and the prolonged interval that follows a sigh, respectively. The distributions remain bimodal with peaks at ∼5 s and ∼10 s after blocking purinergic signaling, indicating that P2/P2Y_1_ receptor blockade does not prevent or modify inspiratory and sigh rhythms. In both conditions, peaks near 1 s reflect the short latency between an inspiratory burst and the subsequent sigh burst (7, 23).

Because the sigh rhythm persists unperturbed after blocking purinergic P2 receptor-mediated signaling, it is unlikely to require gliotransmission, which suggests that astrocytes are not sigh rhythmogenic.

### Sigh and inspiratory rhythms arise from the same neuron population

A previous study reported that sigh-only neurons constitute 5% of the preBötC (15). We monitored Dbx1 preBötC neurons in Dbx1;Ai148 slices (N = 9) by averaging sweeps of their Ca^2+^ transients triggered by discharge of the XII nerve; 208 of 209 Dbx1 preBötC neurons were active during both inspiratory and sigh bursts. We applied a one-sided binomial test to determine the likelihood of results as extreme as ours if sigh-only neurons constitute 5% of the preBötC (*H*_0_: *p* = 0.05): the probability of detecting a single sigh-only neuron given 209 trials is 2.65 × 10^−4^. Therefore, we reject the hypothesis that 5% of preBötC neurons are dedicated to sigh rhythm, which comports with the absence of sigh-only neurons in ref. (16). In summary, our observations provide no evidence to support dichotomous inspiratory and sigh-dedicated neuronal (or glial) populations, but rather, these data suggest both rhythms emerge from the same excitatory *Dbx1*-derived neuronal population already established as inspiratory rhythmogenic (33–35).

### Intracellular Ca^2+^ oscillations produce sigh rhythm

Sigh bursts do not emerge from a voltage-dependent network oscillator (Fig. 1B,C) and Dbx1 preBötC neurons show sigh-related Ca^2+^ transients that do not depend on gliotransmission (Figs. S1 and S2). These observations suggest neuronal Ca^2+^ oscillations produce sigh bursts. A model (13) of the embryonic preBötC posited a similar idea but also depended on *I*_h_. We tested its veracity by applying the selective *I*_h_ blocker ZD7288 (50 μM) to postnatal (not embryonic) slices, which had no effect on sigh frequency (0.87 ± 0.55 min^-1^ in control vs 1.12 ± 0.38 min^-1^ in ZD7288, N = 3, paired t-test p = 0.41). In summary, we found that sigh rhythmogenesis does not depend on *I*_h_ (Fig. S3).

We formulated a mathematical model of the preBötC wherein both inspiratory and sigh rhythms emanate from a single Dbx1 neuron population without an obligatory role for *I*_h_ or chloride-mediated synaptic inhibition (23) (SI Appendix). The model tracks collective network behavior via five state variables for neuronal activity (***a***), synaptic depression (***s***), cellular adaptation (***θ***), and intracellular Ca^2+^ (***c, c***_***tot***_). Peaks in the time series of ***a*** represent inspiratory and sigh bursts (Fig. 2A).

**Figure 2:**
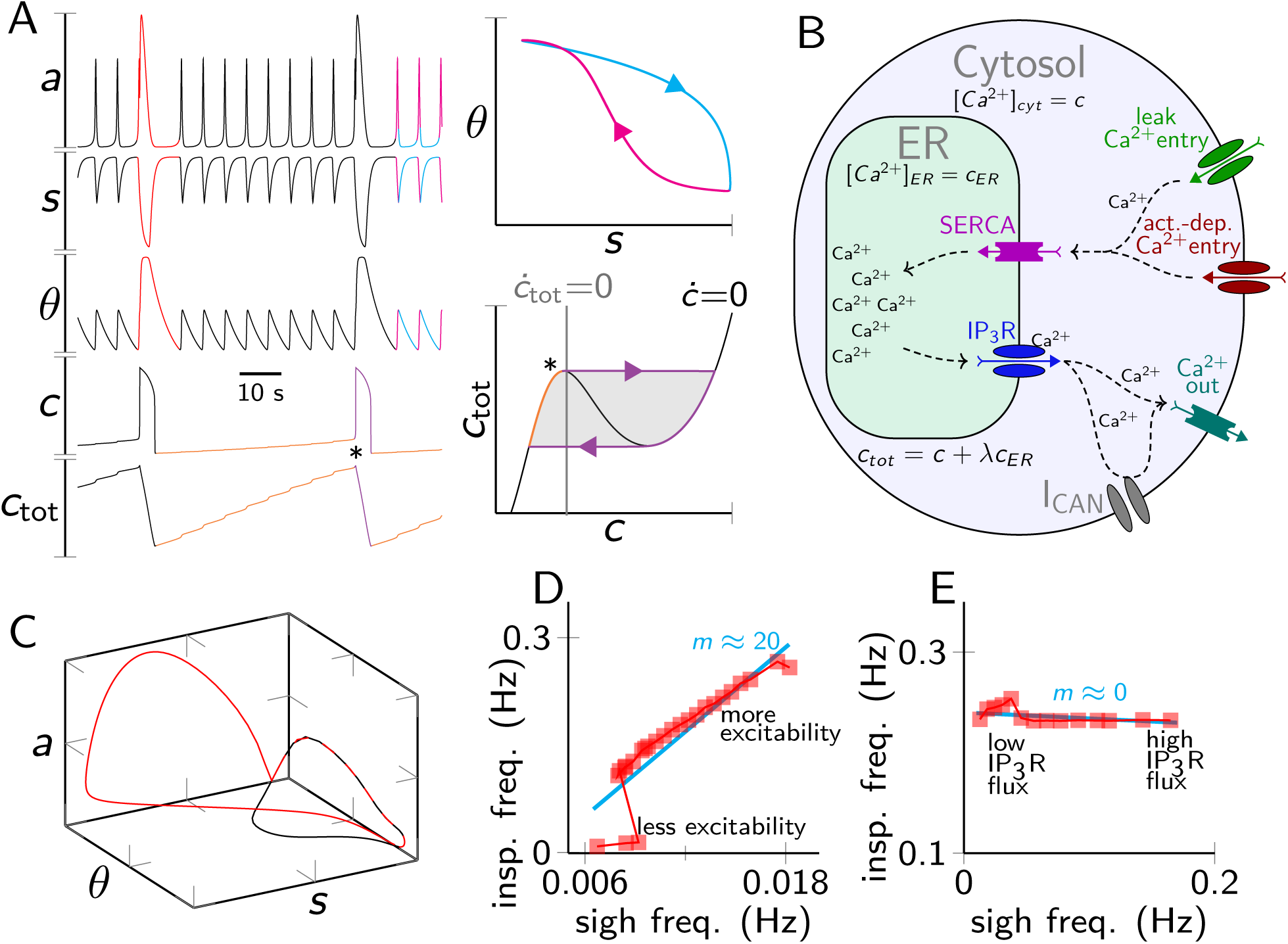
Inspiratory and sigh rhythms in the preBötC model system. (A) At left, time series of state variables (***a, s, θ, c, c***_***tot***_). Top right, inspiratory burst trajectory in (***s, θ***) phase space. Bottom right, sigh trajectory in (***c, c***_***tot***_) phase diagram showing ***c*** and ***c***_***tot***_ nullclines; * marks onset of the sigh burst. (B) Schematic of the Ca^2+^ subsystem showing the critical channels and pumps. Dashed arrows show Ca^2+^ fluxes. (C) Trajectory in (***a, s, θ***) phase space during inspiratory burst events (black) and a sigh event (red). (D,E) Inspiratory and sigh frequencies for different baseline excitabilities for the network (D) or different maximum IP_3_R conductance (***ν***_***ip*3*r***_) (E). Red squares show single simulations; cyan line from linear regression.

The inspiratory subsystem (***a, s, θ***) represents averaged activity and excitability of the preBötC. It is a network oscillator that depends on recurrent excitation, i.e., ***a*** is regenerative during preinspiration (36, 8, 28, 26) (See SI Appendix section 1, Appendix Figs. 1-4). Inspiratory bursts terminate by refractory mechanisms including synaptic depression (***s***) and outward currents recruited by neural activity (***θ***) (24, 10–12). Burst termination is well-illustrated by trajectory in (***s, θ***) phase space where ***s*** decreases (synapses depress) while ***θ*** increases (outward currents activate) (Fig. A, right, magenta). During the interburst interval ***s*** recovers faster than ***θ*** declines (Fig. 2A, right, cyan); the next inspiratory burst occurs when outward currents fully deactivate. This three-variable subsystem captures preBötC burst dynamics (see SI Appendix, section 1) such that it is unnecessary to explicitly model each constituent neuron at present.

Figure 2B shows a schematic of the sigh-rhythmogenic Ca^2+^ subsystem. Inspiratory bursts activate voltage-gated Ca^2+^currents yet cytosolic Ca^2+^ (***c***) increases only minimally because the endoplasmic reticulum (ER) sequesters most of the entering Ca^2+^ via SERCA pumps (37, 38). Total Ca^2+^ (***c***_***tot***_) increases in step with inspiratory rhythm as the ER fills up (Fig. 2A left, orange trace). In the (***c, c***_***tot***_) phase space (Fig. 2A, lower right) the trajectory follows the left (low ***c***) branch of the ***c*** nullcline. When the system reaches its left knee (Fig. 2A, **\***) the replete ER releases Ca^2+^ and the trajectory moves to the right (high ***c***) branch of the ***c*** nullcline (rightward horizontal trajectory, purple arrowhead). The resulting increase in ***c*** evokes Ca^2+^-activated non-specific cation current (*I*_CAN_) to trigger the sigh burst (prominent peaks in the ***a*** time series). During the sigh burst, plasma membrane Ca^2+^ ATPase (PMCA) pumps extrude cytosolic Ca^2+^; ***c*** and ***c***_***tot***_ both decrease (purple trace with down then leftward trajectory) and the system returns to the left (low ***c***) branch of the ***c*** nullcline. Very high ***a*** during a sigh increases ***θ*** beyond its peak value during inspiratory bursts (Fig. 2A,C red traces). ***θ*** takes longer to recover from this elevated level, which explains the post-sigh apnea.

The (***c, c***_***tot***_) system captures Ca^2+^ dynamics within constituent neurons, but can it account for synchronized oscillation throughout the network? We simulated 400 neurons with action potential-generating capabilities and internal Ca^2+^ dynamics as described above. Excitatory synaptic interactions, which are the basis for recurrent excitation, suffice to synchronize the Ca^2+^ oscillations of the constituent neurons (Fig. S4, SI Appendix section 5) such that it is unnecessary to explicitly model each constituent neuron as we test the veracity of our model for explaining the dynamics of the biorhythmic inspiratory-sigh system.

### Inspiratory and sigh rhythms can be separately modulated in the model

We tested model predictions by simulating the change in cellular excitability that results from manipulating [K^+^]_o_ in the aCSF *in vitro*. The relevant model parameter, ***γ***_***a***_, is the input-output function attributable to a leak current that determines how close baseline membrane potentials are to spike threshold. Inspiratory model frequency ranged from quiescence to ∼0.25 Hz as ***γ***_***a***_ varied from 0.1 to 0.4 (akin to varying [K^+^]_o_ 3 to 9 mM) whereas sigh frequency ranged from 0.38 to 1.08 min^−1^ (i.e., 0.006-0.018 Hz, Fig. 2D). Inspiratory rhythm is 20-fold more sensitive to changes in excitability than sigh rhythm, consistent with experiment (compare Figs. 2D and 1C).

We simulated the effects of bombesin-like peptides at the final stage of their Gq-linked signaling cascade (39) by increasing the inositol 1,4,5-trisphosphate receptor (IP_3_R) release rate via parameter ***ν***_***ip*3*r***_. Doing so accelerated sigh frequency from 0.01 to 0.16 Hz without affecting inspiratory frequency (Fig. 2E), which matches the NMB experiments (compare Figs. 2E to 1F). Separate mechanisms and modulation govern inspiratory and sigh rhythms.

### Disrupting SERCA activity reduces sigh frequency

Partially blocking SERCA (i.e., decreasing ***ν***_***SERCA***_ from 60 to 30 s^-1^) counterintuitively increases sigh frequency but decreases sigh magnitude (Fig. 3A). Both effects are a consequence of how SERCA activity influences the ***c***_***tot***_ threshold that compels ER Ca^2+^ release, marked by the left knee of the ***c*** nullcline (Fig. 2A **\***, S5B). Sigh frequency is determined by the time required to refill the ER, which depends on the vertical separation between the knees of the ***c*** nullcline. Reducing ***ν***_***SERCA***_ by 50% decreases their vertical separation and thus speeds-up sigh rhythm (Fig. S5A,B). Further reducing ***ν***_***SERCA***_ by 80% (12 s^-1^) stops sigh rhythmogenesis because the knee of the ***c*** nullcline crosses the vertical ***c***_***tot***_ nullcline, which now intersects the ***c*** nullcline on its positively sloped left branch, i.e., oscillations cease via a Hopf bifurcation (Fig. 3A, S5C).

**Figure 3:**
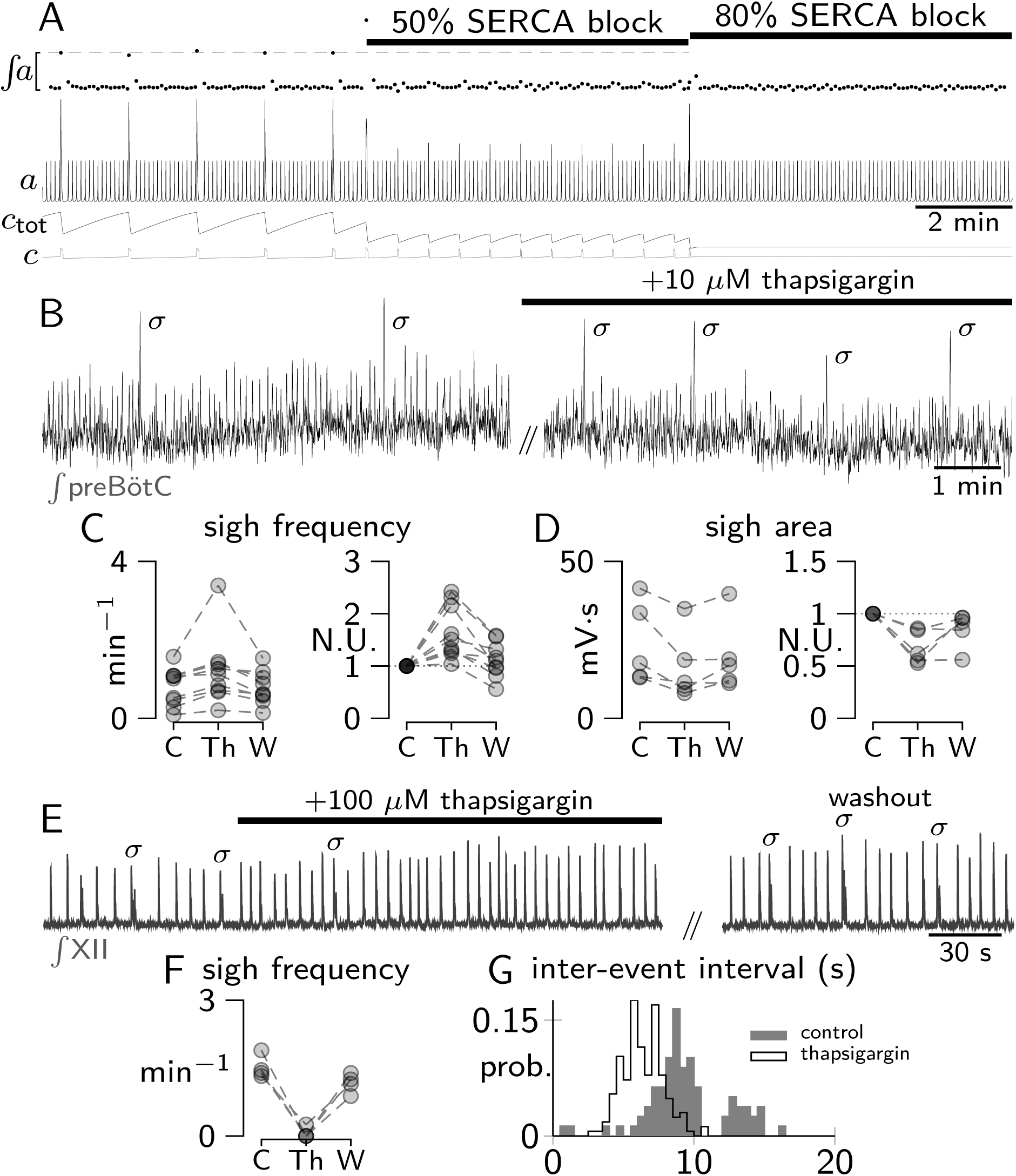
Investigating the role of SERCA pumps in sigh generation. (A) Progressive attenuation of SERCA pump in the model. Filled circles on the ∫ ***a*** axis show area of ***a*** for each burst. Dashed line shows average sigh area during control conditions for reference. (B) preBötC field recordings before and 15 min after application of 10 μM thapsigargin. Sighs are denoted by σ. (C,D) Group data showing average sigh frequency (area) during control (C), thapsigargin (Th), and washout (W) (N = 9 frequency, N = 5 area); data in normalized units (NU) also shown. (E) XII nerve recording during 100 μM thapsigargin application and washout. (F) Sigh frequency during control (C), thapsigargin (Th), and washout (W) (N=4). (G) Inter-event interval distributions.

Sigh frequency in slices can be determined from preBötC field recordings or via XII motor output. However, sigh magnitude can only be accurately measured via preBötC field recordings because XII output is filtered by premotor neurons postsynaptic of the preBötC core. We bath applied 10 μM thapsigargin to partially block SERCA pumps. Sigh frequency increased 0.8 ± 0.48 min^-1^ in control vs. 1.23 ± 0.90 min^-1^ in thapsigargin (paired t-test, p = 0.04, N = 9) (Fig. 3B,C). Sigh burst magnitude decreased by 32% (24 ± 13 mV-s in control vs. 16 ± 11 mV-s in thapsigargin, paired t-test, p = 0.029, N = 5) (Fig. 3B,D). The effect on sigh frequency was reversible (0.8 ± 0.42 min^-1^); sigh magnitude recovered in three out of the five field recordings (19 ± 11 mV-s) (Fig. 3C,D). These experimental results matched the model predictions for partial SERCA blockade.

Next, we fully blocked SERCA pumps by injecting a high dose of thapsigargin (100 μM) bilaterally into the preBötC. We microinjected into the preBötC to avoid widespread blockade of SERCA pumps following bath application. Local microinjection is important in this context because neurons outside of preBötC that are retained in slices, like the raphé obscurus, are tonically active and help maintain preBötC excitability (40). Widespread SERCA blockade via bath application of a high concentration of thapsigargin could impact the preBötC rhythmogenic network via indirect effects. The drawback is that local injection precludes simultaneous local field potential recording. Therefore, we assessed sigh rhythm following 100 μM thapsigargin solely via XII output and thus only analyzed sigh frequency.

To establish that our injection micropipettes correctly targeted the preBötC we first injected bolus of 25 mM [K^+^]_o_ aCSF into the preBötC bilaterally. Potassium rapidly and reversibly increased inspiratory frequency whereas 1% DMSO aCSF, the vehicle, had no effect (Fig. S6). Thapsigargin (100 μM) stopped the sigh rhythm (N = 3) or decreased it by 81% to 0.25 min^-1^ (N = 1) (Fig. 3E,F), which recovered in washout (from 1.52 ± 0.25 min^-1^ in control to 1.17 ± 0.21 min^-1^ in washout).

We further measured inter-event intervals as a metric of sigh rhythm. During control conditions the inter-event interval distribution is bimodal, with a peak at ∼8 s representing the inspiratory intervals and a peak at ∼13 s representing the post-sigh apneas (Fig. 3G). The inter-event interval histogram became monophasic in the presence of 100 μM thapsigargin; its sole peak at ∼6 s represents inspiratory intervals; the lack of another peak at >6 s indicates the cessation of sigh rhythm. These experimental results matched the model predictions for ≥80% blockade of SERCA pumps.

### Blocking IP_3_Rs diminishes sigh frequency

Attenuating IP_3_R-mediated Ca^2+^-induced Ca^2+^ release rate (***ν***_***ip*3*r***_) at first decelerates and then stops the sigh rhythm (Fig. 4A). Decreasing ***ν***_***ip*3*r***_ ≤40% elevates the critical ***c***_***tot***_ that compels ER Ca^2+^ release (Fig. 2A **\***), enhancing the vertical separation between the knees of the ***c*** nullcline and slowing-down the sigh rhythm (Fig. 4A, S7A,B). ***ν***_***ip*3*r***_ reduction did not affect sigh burst area as indicated by the ∫ *a* in Fig. 4A. Decreasing ***ν***_***ip*3*r***_ ≥50% raises the left knee high enough to cross the vertical ***c***_***tot***_ nullcline, which now intersects the ***c*** nullcline on its positively sloped left branch, i.e., oscillations cease via a Hopf bifurcation (Fig. 4A, S7C).

**Figure 4:**
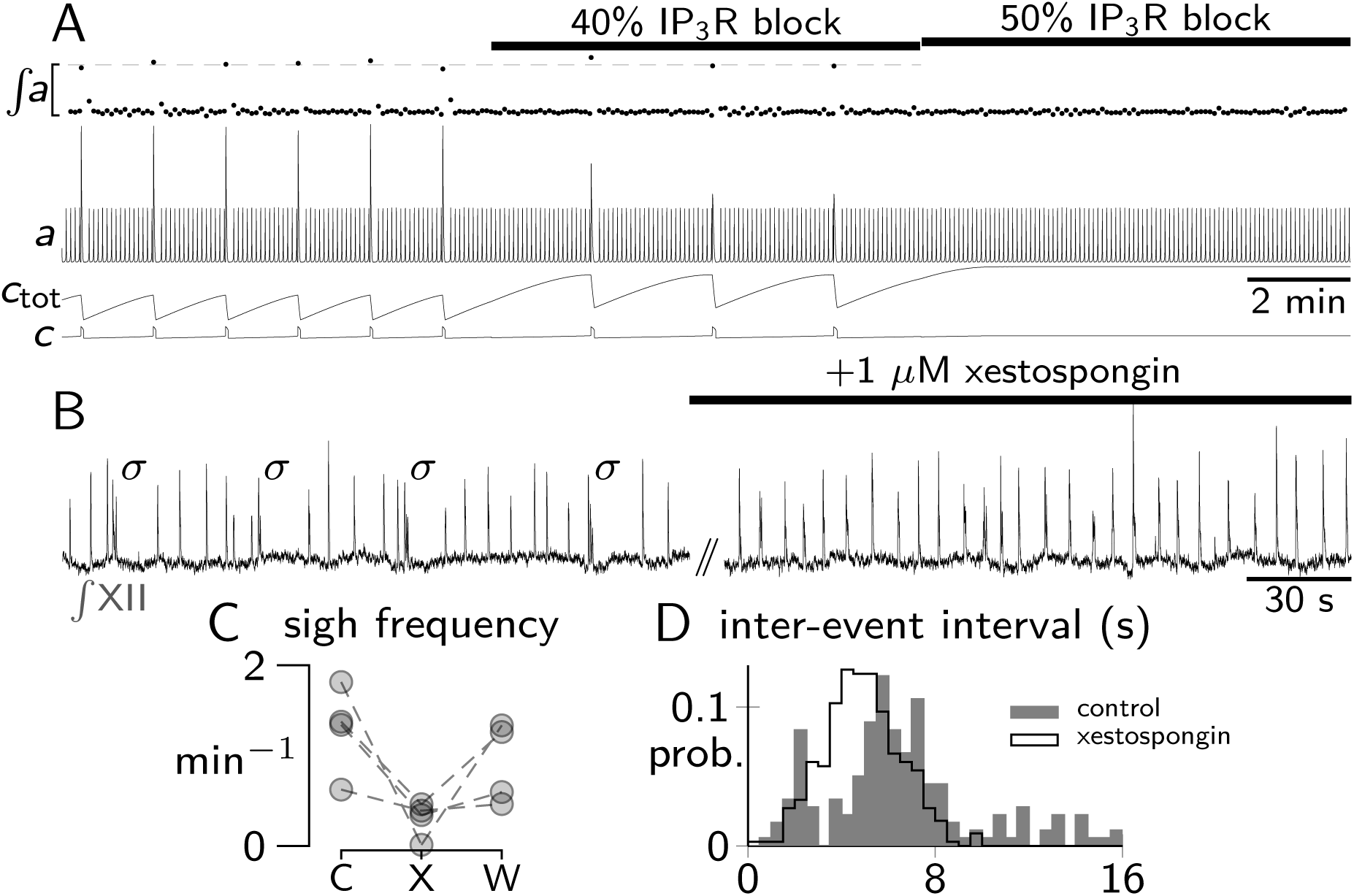
Investigating the role of IP_3_Rs in sigh generation. (A) Progressive IP_3_R blockade in the model. Filled circles on the ∫ ***a*** axis show area of ***a*** for each burst. Dashed grey line shows average sigh area during control conditions for reference. (B) XII nerve recordings before and during application of 1 μM xestospongin. Sighs are denoted by σ. (C) Sigh frequency during control (C), xestospongin (X) application, and washout (W) (N=4). (D) Inter-event interval distributions.

We tested this model prediction in slices by attenuating IP_3_Rs using xestospongin (41, 42), injected bilaterally into the preBötC. It was impracticable to calibrate xestospongin dose to mimic 40% vs. 50% attenuation of ***ν***_***ip*3*r***_. Xestospongin (1 μM) decreased sigh frequency (1.29 ± 0.49 min^-1^ in control vs. 0.3 ± 0.2 min^-1^ in xestospongin, N = 3) or stopped it altogether (N = 1), in broad agreement with model predictions (Fig. 4B,C). The effect was reversible; sigh frequency returned to 0.91 ± 0.48 min^-1^.

We further measured inter-event intervals as a metric of sigh rhythm. During control conditions the inter-event interval distribution is bimodal, with a peak at ∼6 s representing the inspiratory intervals and a broad peak at ∼12 s representing the post-sigh apneas (Fig. 4D). Xestospongin caused the peak ∼12 s disappear, which indicates the cessation sigh rhythm.

## Discussion

Eupnea and sigh rhythms both come from the preBötC (7) but from which cell population(s)? Furthermore, could the slower rhythm, lacking voltage dependence, depend on glia? How are the rhythms coupled? Here we resolve these questions by showing that both rhythms emanate from one neuronal population in the preBötC postnatally with no obligatory role for purinergic gliotransmission. Membrane properties like *I*_h_ and synaptic inhibition (ref. 23) are not required for rhythmogenesis or coupling, respectively.

We conclude that glia and gliotransmission are not mandatory for sigh rhythmogenesis because attenuating purinergic P2Y signaling, the dominant means by which astrocytes interact with preBötC neurons (30–32), did not preclude or modify sigh rhythmogenesis. A recent report in preprint form showed that blocking gliotransmission at P2Y_1_ receptors stops the sigh rhythm *in vitro* but does not stop sighs in adult mice (17). Our conclusion is not incompatible – purinergic gliotransmission is ultimately unnecessary for sigh behavior – but our data *in vitro* are incongruous. The disparity may be attributable to which populations were monitored. We recorded Dbx1 neurons in the preBötC core from the rostral surface of 500 μm-thick slices. In contrast, ref. 17 performed field recordings from the caudal surface of ∼600 μm-thick slices, in which the recording site is ∼200 μm caudal to the preBötC in an area containing phrenic premotor neurons (6). Therefore, cessation of sigh rhythm suggests that P2/P2Y_1_ receptor-mediated signaling might be critical for premotor transmission of sigh bursts while remaining dispensable from the standpoint of sigh rhythmogenesis.

Our conclusion that glia are not sigh rhythmogenic assumes that signaling is purely purinergic. However, astrocytes in the trigeminal system facilitate oromotor rhythmogenesis via paracrine transmission (43). There, S100*β* secretion neither generates nor synchronizes oscillations but rather modulates *I*_NaP_-mediated bursting-pacemaker properties in rhythmogenic trigeminal interneurons. This mechanism is highly unlikely to apply to sigh rhythmogenesis because *I*_NaP_ bursting-pacemaker neurons oscillate much faster than the sigh rhythm and are not inspiratory rhythmogenic in the preBötC (18–21).

One report found 5% of preBötC neurons were active only during sighs and postulated them as sigh rhythmogenic (15). A contemporary report found the opposite, namely that all preBötC neurons participated in both sighs and inspiratory bursts (16). We monitored Dbx1 preBötC neurons via photonic imaging; the vast majority were bi-rhythmically active. We conclude that inspiratory and sigh rhythms come from a single population of Dbx1 preBötC neurons.

One neuron population produces two rhythms by engaging separate cellular tool kits. The faster inspiratory rhythm employs recurrent synaptic interactions (9, 27, 28, 33, 34); it is a canonical network oscillator (1) whose bursts emerge on the order of seconds. Nevertheless, each constituent neuron hosts an intracellular signaling system linked to plasma membrane Ca^2+^ flux as well as ER Ca^2+^ storage and release mechanisms. That system can produce biochemical oscillations much slower than inspiration and trigger neural bursts substantially larger than inspiration via Ca^2+^-induced Ca^2+^ release that evokes the burst-generating inward current, *I*_CAN_. The two mechanisms can comfortably coexist in preBötC neurons because the underlying network oscillator and a biochemical oscillator are fundamentally different and can be separately regulated by manipulating either membrane excitability (inspiration) or Gq-mediated intracellular signaling (sigh).

The two oscillators have a second point of intersection: plasma membrane Ca^2+^ influx. Network activity increases Ca^2+^ influx leading to a modest increase Ca^2+^ oscillation frequency. This confers (minimal) voltage dependence on the sigh oscillation, which is approximately 5% as sensitive as inspiratory rhythm to changes in membrane excitability (see Figs. 1B,C and 2D). Activity dependent Ca^2+^ influx may also promote synchronization of intracellular Ca^2+^ oscillations across the neural population. In the spiking model, ionotropic synaptic signaling can maintain synchronized intracellular Ca^2+^ oscillations underlying the sigh rhythm (Fig. S4 and SI Appendix Fig. 12). In the biological system, additional mechanisms are available including metabotropic glutamatergic transmission (21) as well paracrine signaling mechanisms yet to be identified.

The evolution of the compliant lung (one that expands and contracts) in mammals introduced the physiological need for regularly timed large-volume breaths to reinflate collapsed or collapsing alveoli. As the breathing system evolved from fish to reptiles it produced an inexorable inspiratory oscillator but not a sigh rhythm. As the physiological need arose, rather than recruit a new brain region for sighs, we posit that the mammalian nervous system adapted the inspiratory rhythmogenic neurons to produce a low frequency, large amplitude oscillation that depended on a separate intracellular tool kit in the same constituent Dbx1 preBötC neurons.

This work demonstrates the operation of a bi-rhythmic canonical CPG system with relevance to health and physiology. Its dynamics can be understood via a low-dimensional dynamical system with fast and slow time scales.

## Materials and methods

### Ethical approval and animal use

The Institutional Animal Care and Use Committee at William & Mary approved these protocols, which conform to the policies of the Office of Laboratory Animal Welfare (National Institutes of Health, Bethesda, MD, USA) as well as the guidelines of the National Research Council (44). Mice (described below) were maintained on a 12-hour light / 12-hour dark cycle at 23° C and were fed *ad libitum* with free access to water. The mice are provided with several forms of enrichment including opaque igloo shelters, wood blocks, and nest materials.

Multi-photon experiments employed Cre-driver mice generated by inserting an *IRES-CRE-pGK-Hygro* cassette in the 3’ untranslated region of the *Developing brain homeobox 1* (i.e., *Dbx1*) gene, which we refer to as *Dbx1*^*Cre*^ mice (45) (IMSR Cat# EM:01924, RRID:IMSR_EM:01924). We crossed female *Dbx1*^*Cre*^ mice with males from a reporter strain featuring Cre-dependent expression of the fluorescent Ca^2+^ indicator GCaMP6f dubbed Ai148 by the Allen Institute (46) (IMSR Cat# JAX:030328, RRID:IMSR_JAX:030328). Their offspring, Dbx1;Ai148 mice, express GCaMP6f in *Dbx1*-derived cells, the majority of which are neurons (47).

### Breathing-related measurements in vitro

Mouse pups of both sexes were anesthetized by hypothermia and killed by thoracic transection at postnatal day 0 to 4. Neuraxes were removed in artificial cerebrospinal fluid (aCSF) containing (in mM): 124 NaCl, 3 KCl, 1.5 CaCl_2_, 1 MgSO_4_, 25 NaHCO_3_, 0.5 NaH_2_PO_4_, and 30 dextrose equilibrated with 95% O_2_-5% CO_2_, pH 7.4. Isolated neuraxes were glued to an agar block and then cut in the transverse plane to obtain a single 500-μm-thick slice that exposed the preBötC at its rostral face. Atlases for wild-type and Dbx1 reporter mice show that the loop of the inferior olive and the semi-compact division of the nucleus ambiguus collocate with the preBötC during early postnatal development (48, 49). Slices were then perfused with aCSF at 28º C in a recording chamber below a fixed-stage microscope.

We elevated extracellular K^+^ concentration ([K^+^]_o_) to 9 mM to increase preBötC excitability (50). Inspiratory-related motor output was recorded from the hypoglossal (XII) nerve rootlets, which are captured in transverse slices along with the XII motoneurons and their axon projections to the nerve rootlets, using suction electrodes and a differential amplifier. We obtained field potential recordings by forming a seal over the preBötC with a suction electrode at the rostral slice surface. Amplifier gain was set at 1000. Signals were acquired digitally at 1 kHz while low-pass filtering at 300 Hz. XII and preBötC bursts were full-wave rectified and smoothed for display and quantitative analyses of burst events.

To apply drugs locally in the preBötC we fabricated micropipettes from borosilicate glass (OD: 1.5 mm, ID: 0.86 mm) and filled them with either thapsigargin or xestospongin (see below). Two pipettes were inserted 200 μm deep into the preBötC on both sides of slice preparations. We microinjected the drugs using 7-9 psi pressure pulses lasting 8 ms in duration, delivered at a frequency of 5-7 Hz. Pipettes for local drug application in the preBötC precluded simultaneous local field recordings; we monitored preBötC activity in those experiments only via XII nerve recordings.

### Multi-photon imaging

We imaged cytosolic Ca^2+^ concentration in neurons contained in slices from Dbx1;Ai148 mice using a multi-photon laser-scanning confocal microscope (Thorlabs, Newton, NJ) equipped with a Nikon water immersion objective (20x, 1.0 numerical aperture). Illumination was provided by an ultrafast tunable laser with a power output of 950 mW at 940 nm, 80-MHz pulse frequency, and 100-fs pulse duration (Coherent Chameleon Discovery, Santa Clara, CA). We scanned Dbx1;Ai148 mouse slices over the preBötC and collected time series images using a non-descanned photomultiplier tube detector at 15 Hz. Each frame reflects one-way raster scans with a resolution of 256 × 256 pixels (116 × 116 μm). Fluorescence data were collected using Thorlabs LS 4.1 software and then analyzed using MATLAB 2021a (MathWorks, Natick, MA, RRID:SCR_001622).

First, we calculated the average fluorescence intensity for all pixels in each frame of the time series. The mean fluorescence intensity was used as an index of overall network activity during the time series. The bursts of fluorescence intensity were periodic and the cycle periods were normally distributed. We use the 95% confidence interval (CI) of cycle periods to define the high frequency (short cycle period) and low frequency (long cycle period) limits of a window in frequency space. Next, our script performs a fast Fourier transform on the time series for each pixel. The maximum power from the previously defined window in frequency space is mapped to the corresponding pixel in a new, processed two-dimensional image.

We calculate the mean and standard deviation of the power from each pixel in the new processed image (Fig. S1A). Rhythmically active pixels will have power far greater than the average. Therefore, all pixels with intensity less than mean + 2**\***SD are set to zero. The remaining contiguous pixel sets, whose area exceeds 8 μm^2^, are retained as ROIs. The Ca^2+^ fluoresce changes within those ROIs, obtained from the original time series, are reported using the equation 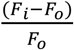, i.e., 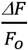, where *F*_*i*_ is the instantaneous average fluorescence intensity for all pixels within a given ROI and *F*_*o*_ is the average fluorescence intensity of all pixels within that same ROI averaged over the entire time series.

### Inspiratory burst and sigh burst detection

We distinguished a sigh burst from an inspiratory burst in the preBötC field recordings by measuring burst area and the duration of the interval between the putative sigh burst and the subsequent inspiratory burst. Sigh bursts are typically ≥2x larger in area than inspiratory bursts. Additionally, a prolonged interval between the putative sigh and the following inspiratory burst, typically 1.3x the average inter-event interval for inspiratory bursts, confirms that the event in question is a sigh burst.

### Pharmacology

We employed the following drugs to block neuron-glia signaling: PPADS (50 μM), suramin (50 μM), TNP-ATP (10 μM), MRS2179 (10 μM), and MRS2578 (10 μM). We used thapsigargin (10-100 μM) and xestospongin C (1 μM) to interrogate intracellular Ca^2+^ sequestration and release. We employed ZD7288 (50 μM) to block *I*_h_. Thapsigargin, xestospongin, and ZD7288 were dissolved in dimethyl sulfoxide (DMSO) to generate stock solutions. Final concentration of DMSO in aCSF never exceeded 1% by volume. All drugs were obtained from Millipore Sigma (Burlington, MA).

### Numerical simulations and data analysis

We used MATLAB 2021a and XPPAUT software to simulate and analyze ordinary differential equation models. Numerical integration was performed using Euler’s method with a time step of 0.01 ms in MATLAB. XPPAUT was used with default solver settings. The SI Appendix describes the modeling work in detail. The code and equations are in the public repository on ModelDB (Accession No. 267252).

Error bars show standard deviation. P-values for regression statistics are calculated using the *F*-test.

## ACKNOWLEDGEMENTS AND FUNDING SOURCES

This work was supported by the National Institutes of Health grants R01-HL104127 (PI: CA Del Negro) and AT010816 (PIs: Conradi Smith and Del Negro), and National Science Foundation grant DMS 1951646 (PI: Conradi Smith).

## Notes

### Competing Interest Statement

The authors have declared no competing interest.

