## Supplemental Information and Appendix for "Sigh breathing rhythm depends on intracellular calcium oscillations in a population of inspiratory rhythmogenic preBötzinger complex neurons in mice"

##### This PDF file includes:

Figs. S1 to S6
Appendix: Modeling of inspiratory and sigh rhythms

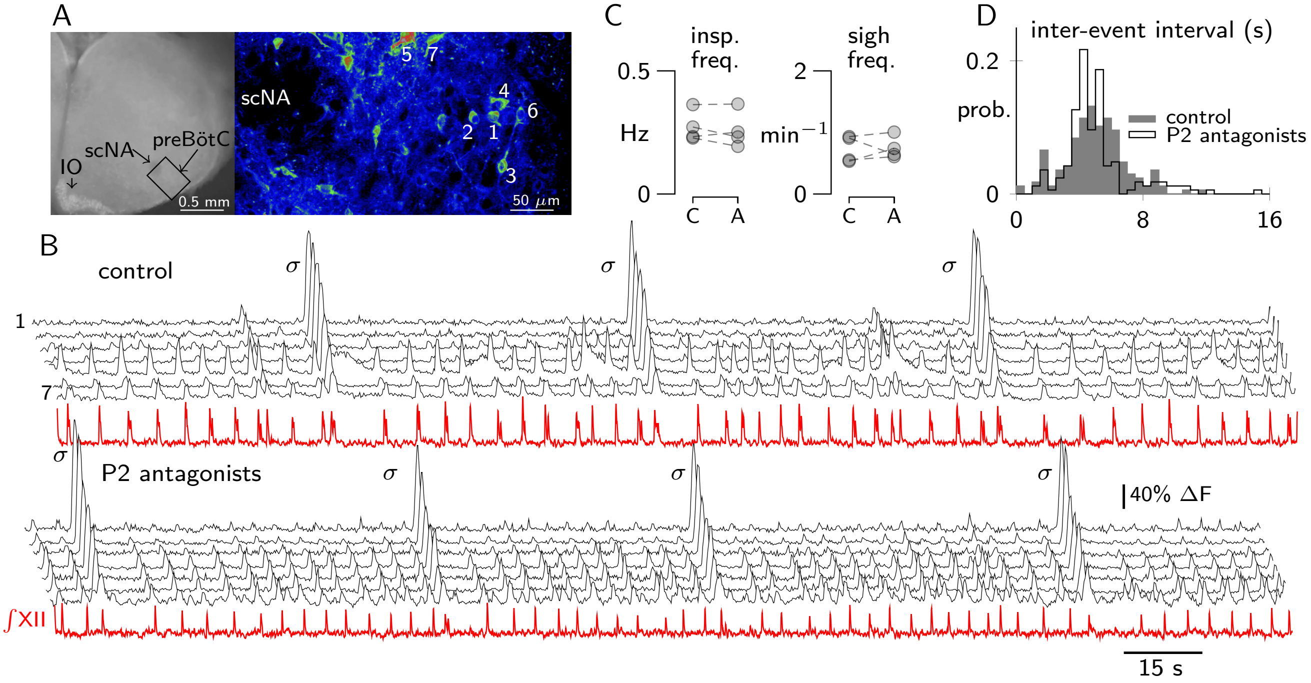


**Figure S1.** Effects of a cocktail of P2 receptor antagonists on inspiratory and sigh rhythms. (A) In vitro slice preparation (left). Characteristic sites indicated include the preBötzinger complex (preBötC), semi-compact nucleus ambiguus (scNA), and the inferior olivary nucleus (IO). Bounding box centered on the preBötC is magnified and rotated 45° at right. Pseudocolor pixel intensity indicates inspiratory rhythmicity obtained by frequency domain analyses (see Materials and Methods). Numerals correspond to regions of interest, i.e., Dbx1 preBötC neurons, rhythmically fluorescing in sync with the inspiratory rhythm displayed in B. (B) $Ca^{2+}$ fluorescence changes in Dbx1 preBötC neurons with XII motor output (red). Unmarked bursts are inspiratory; $\sigma$ indicates sigh bursts. The cocktail of P2 receptor antagonists includes PPADS (50 $\mu$M), Suramin (50 $\mu$M), TNP-ATP (10 $\mu$M), MRS2179 (10 $\mu$M), and MRS2578 (10 $\mu$M). (C) Mean inspiratory and sigh frequency for each slice tested in control (C) and P2 antagonists (A) conditions. (D) Distribution of inter-event intervals for all cycles in all slices tested in control and P2 antagonists conditions.

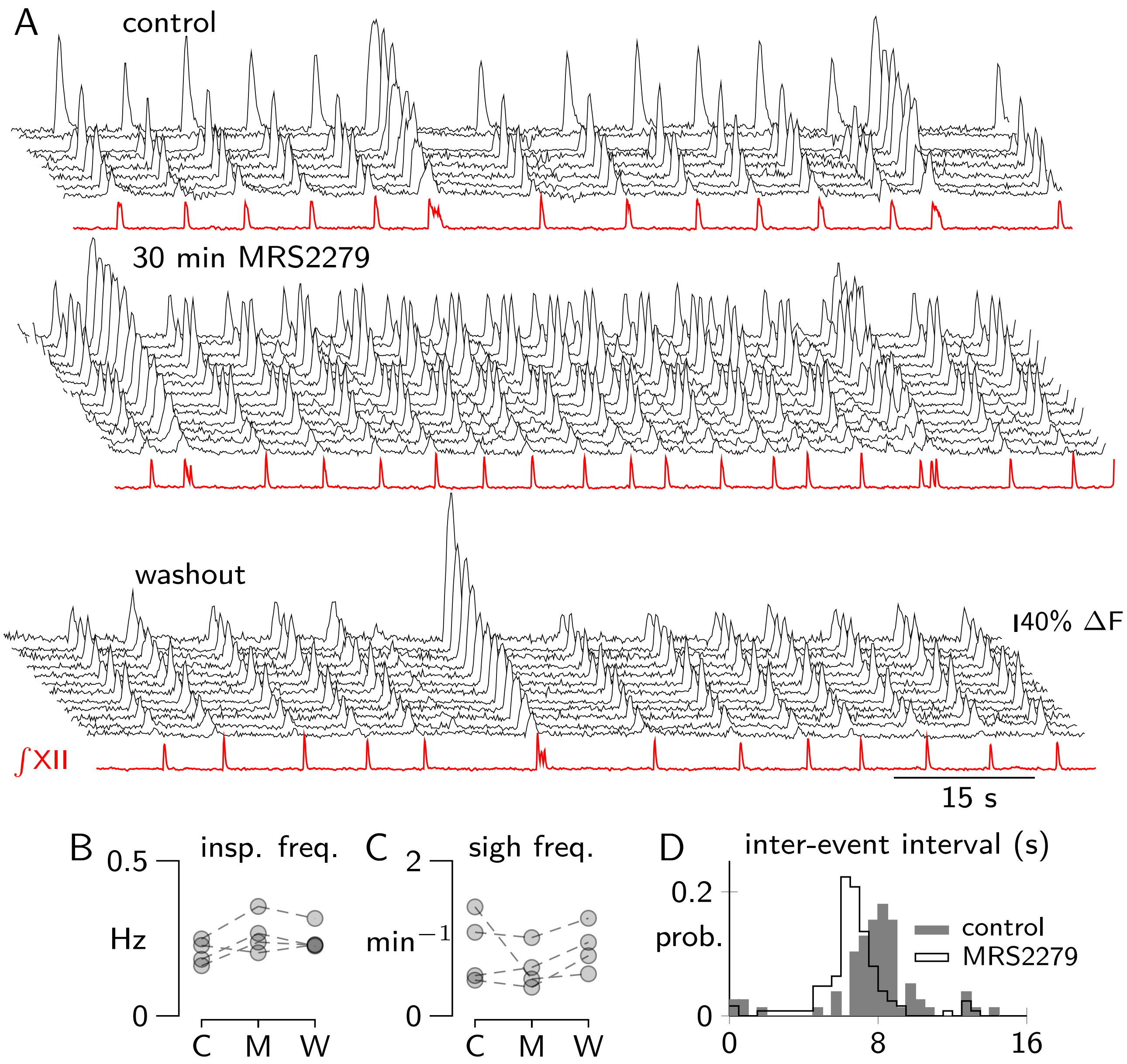


**Figure S2.** Effect of specific P2Y1 receptor antagonist MRS 2279 on inspiratory and sigh rhythms. (A) $Ca^{2+}$ fluorescence changes in Dbx1 preBötC neurons with XII motor output (red). MRS 2279 was applied at 20 $\mu$M. (B, C) Mean inspiratory (B) and sigh (C) frequency for each slice tested in control (C), MRS 2279 (M), and washout (W) conditions. (D) Distribution of inter-event intervals for all cycles in all slices tested in control and MRS 2279 conditions.

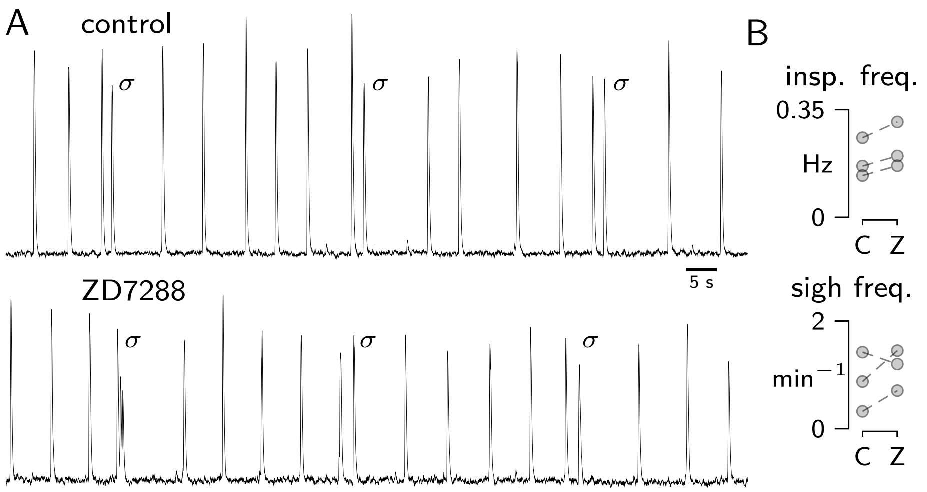


**Figure S3.** Effects of ZD 7288 on inspiratory and sigh rhythms. (A) XII output from a slice preparation. Unmarked bursts are inspiratory; $\sigma$ indicates sigh bursts. ZD 7288 was applied at 50 $\mu$M. (B) Mean inspiratory and sigh frequency for each slice tested in control (C) and ZD 7288 (Z) conditions.

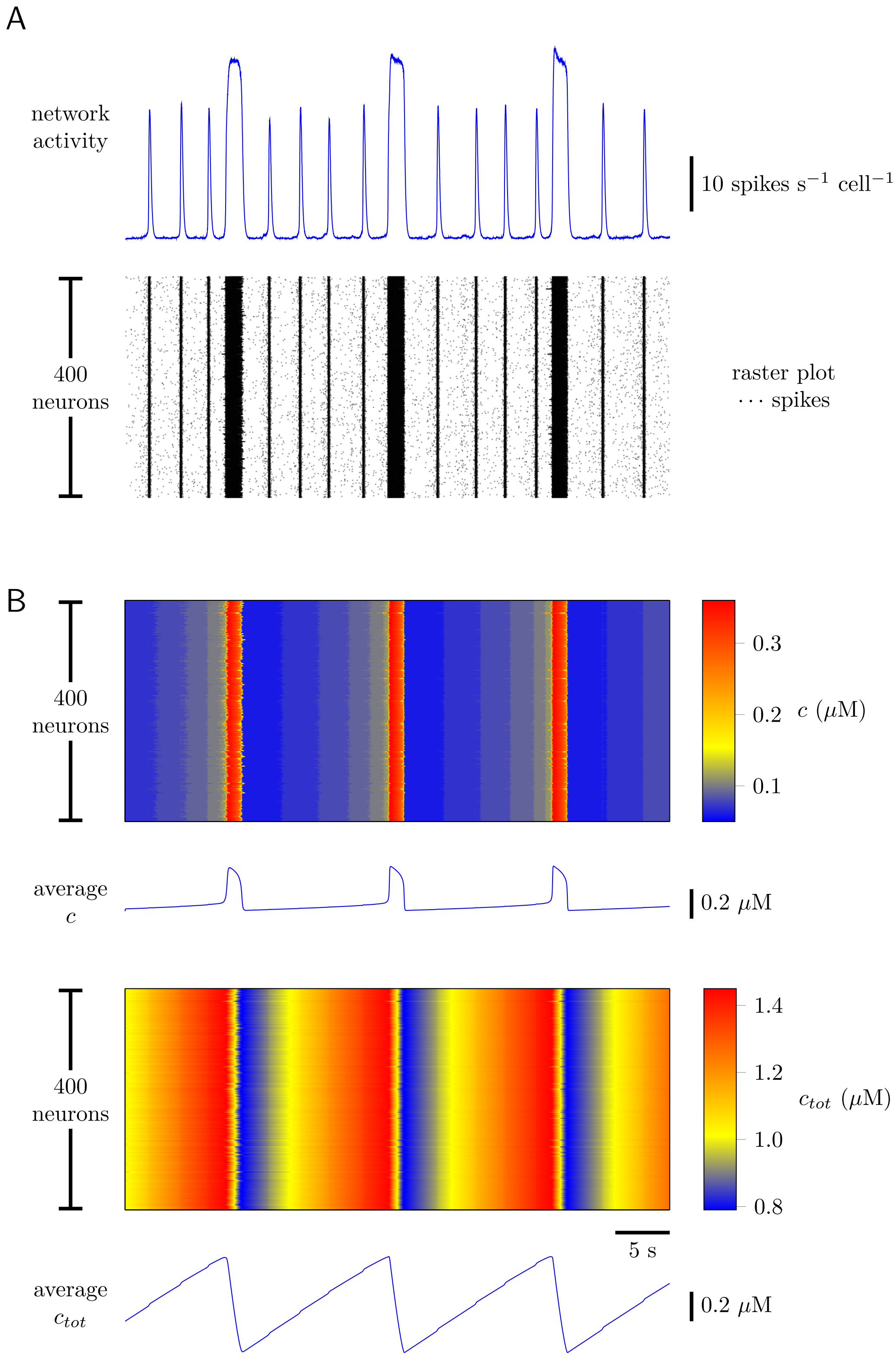


**Figure S4.** Network activity and dynamics of intracellular [Ca$^{2+}$] for a spiking model of eupnea and sigh rhythmogenesis. The network structure is Erdös-Rényi-type with 400 neurons and 6.5% probability that any given neuron is postsynaptic to any other. (A) Network activity and raster plot. (B) Cytosolic [Ca$^{2+}$] ($c$) and total [Ca$^{2+}$] ($c_{tot}$) are highly synchronized across the 400 neurons. See Section 5 of SI Appendix for model description, equations, and parameters.

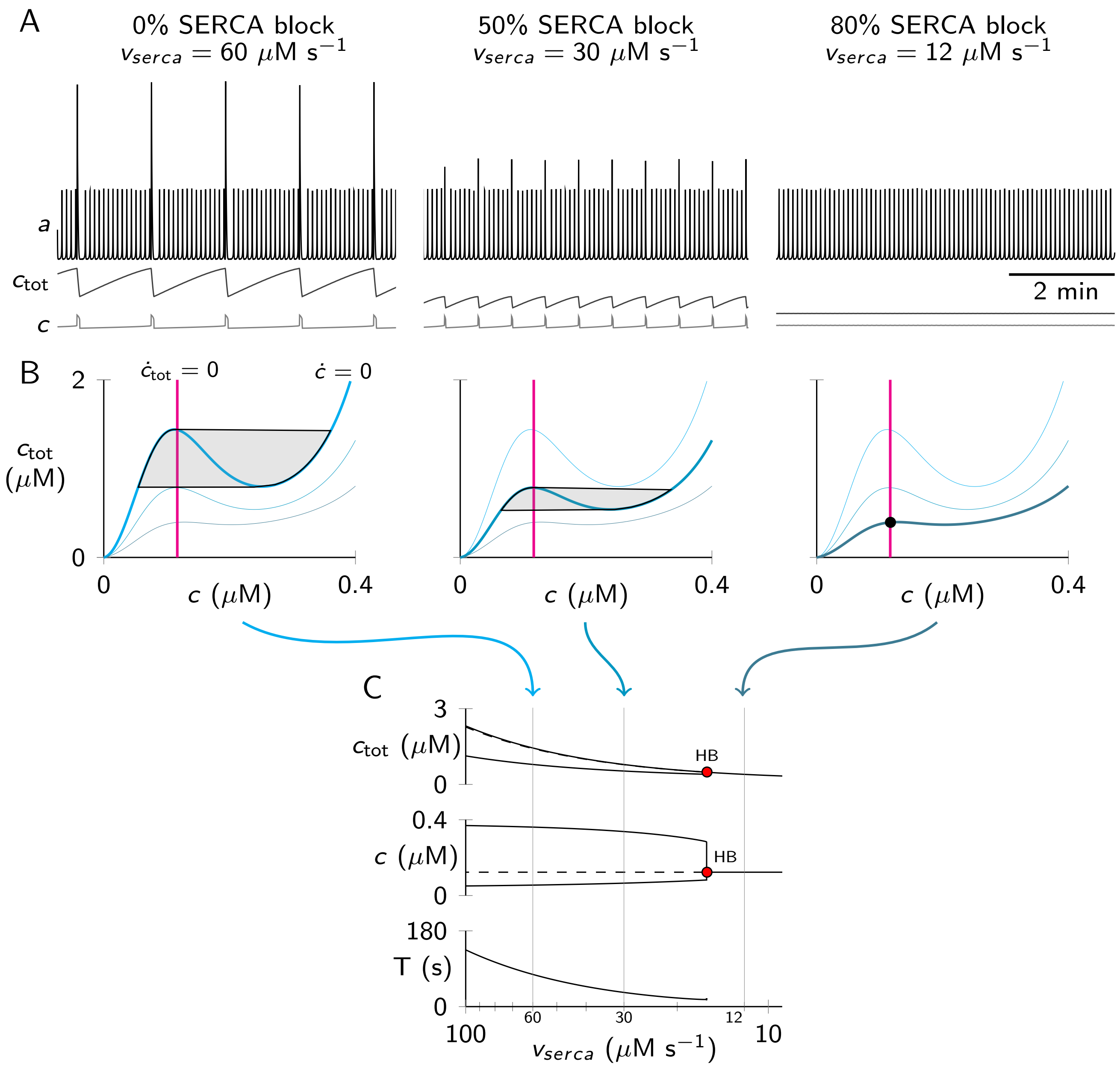


**Figure S5.** Effects of SERCA blockade in the model. (A) Time series showing network activity ($a$), cytosolic $Ca^{2+}$ ($c$), and total $Ca^{2+}$ ($c_{tot}$) in control and after reducing vserca by 50% and then 80%. (B) Model behavior in the $c$-$c_{tot}$ phase plane for corresponding values of $v_{serca}$. Cyan shows the $c_{tot}$ nullcline; magenta shows the $c$ nullcline. Shaded areas indicate limit cycles (direction clockwise); shaded point indicates a stable steady state (no oscillations). (C) Bifurcation diagrams for $c_{tot}$, $c$, and cycle period ($T$) versus the parameter $v_{serca}$ plotted logarithmically in descending order from 100 to 9 $\mu M\,s^{-1}$. HB indicates a Hopf bifurcation point. $v_{serca}$ values from the time series (A) and phase planes (B) are indicated on the abscissa.

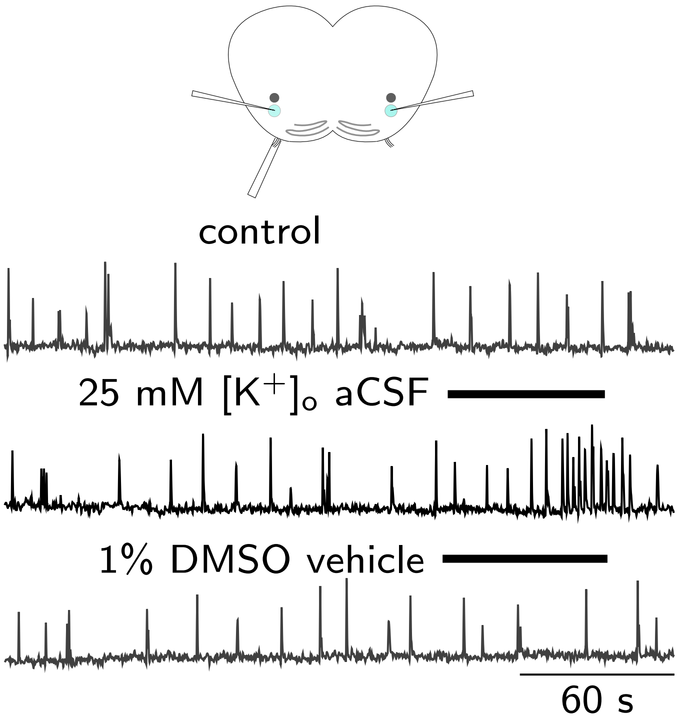


**Figure S6.** Validation of bilateral microinjection methods. Cartoon shows a slice preparation with XII recording pipette and two injection pipettes in the preBötC bilaterally. Robust yet transient frequency response indicates that the pipettes accurately target the preBötC. Injection of the vehicle dimethylsulfoxide (DMSO) is without effect.

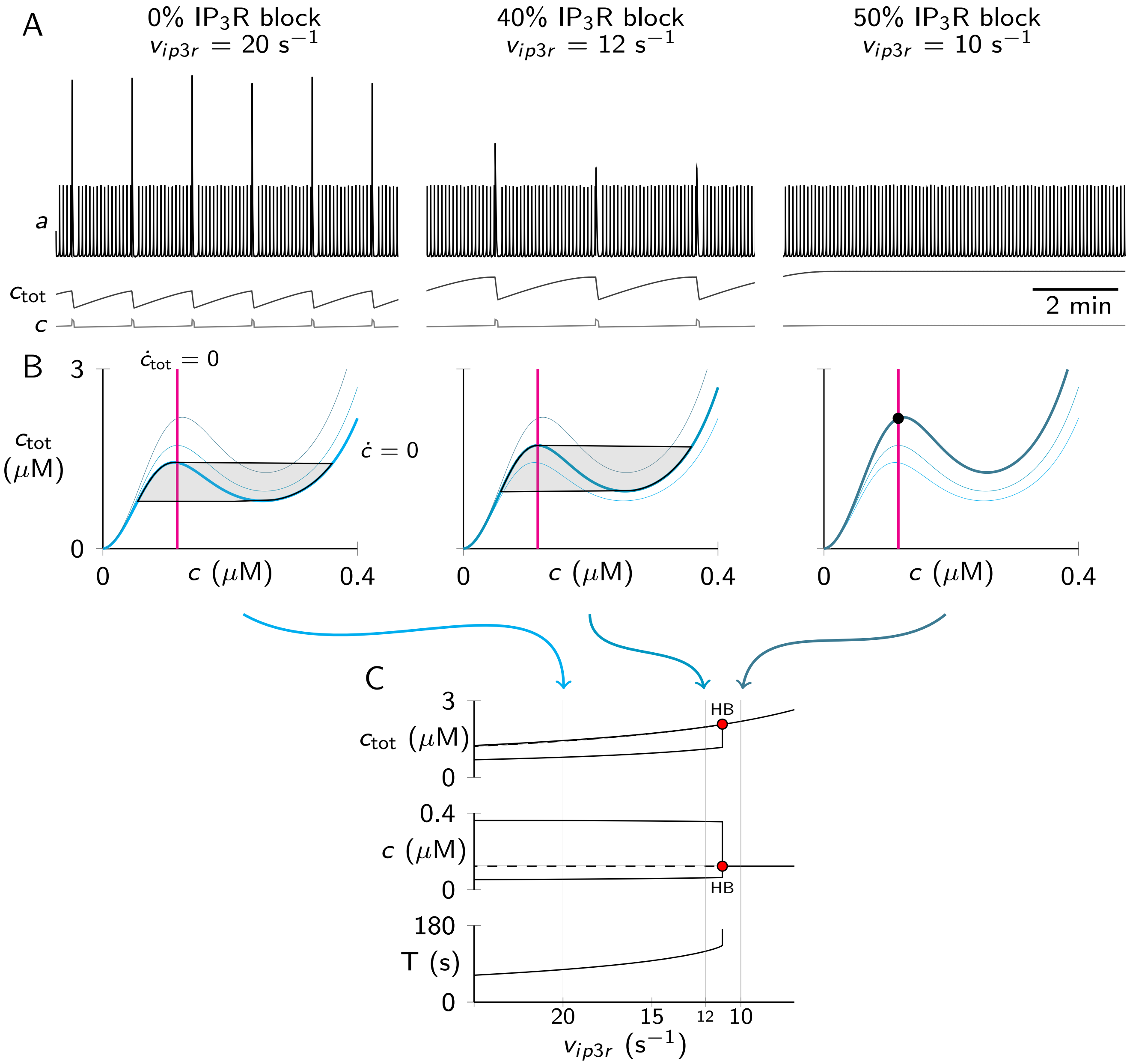


**Figure S7.** Effects of $IP_3R$ blockade in the model. (A) Time series showing network activity ($a$), cytosolic $Ca^{2+}$ ($c$), and total $Ca^{2+}$ ($c_{tot}$) in control and after blocking $IP_3Rs$ by 40% ($v_{ip3r}$ reduced by 60%) and then blocking $IP_3Rs$ 50% ($v_{ip3r}$ reduced by 50%). (B) Model behavior in the $c$-$c_{tot}$ phase plane for corresponding values of $v_{ip3r}$. Cyan shows the $c_{tot}$ nullcline; magenta shows the $c$ nullcline. Shaded areas indicate limit cycles (direction clockwise); shaded point indicates a stable steady state (no oscillations). (C) Bifurcation diagrams for $c_{tot}$, $c$, and cycle period ($T$) versus the parameter $v_{ip3r}$ plotted in descending order from 25 to 8 $s^{-1}$. HB indicates a Hopf bifurcation point. $v_{ip3r}$ values from the time series (A) and phase planes (B) are indicated on the abscissa.

### Appendix: Modeling of inspiratory and sigh rhythms

The model of inspiratory and sigh rhythmogenesis (Appendix Fig. 1 below and Fig. 2 of the main text) is an ordinary differential equation (ODE) system with five dynamical variables—three for the inspiratory activity of the preBötzinger complex (preBötC) neuronal network ($a$, $s$, $\theta$) and two for the dynamics of intracellular calcium ($Ca^{2+}$) in a representative preBötC neuron ($c$, $c_{tot}$). The differential equations for the full model are

$$\tau_a \frac{da}{dt} = a_\infty(w \cdot s \cdot a - \theta, c) - a \quad [1]$$

$$\tau_s \frac{ds}{dt} = s_\infty(a) - s \quad [2]$$

$$\tau_\theta(a) \frac{d\theta}{dt} = \theta_\infty(a) - \theta \quad [3]$$

$$\frac{dc}{dt} = \underbrace{[v_{ip3r} f_{open}(c) + v_{leak}]\,[c_{er}(c, c_{tot}) - c]}_{j_{rel}} - \underbrace{\frac{v_{serca} c^2}{\kappa_{serca}^2 + c^2}}_{j_{serca}} + \underbrace{j_0 + j_a a}_{j_{in}} - \underbrace{\frac{v_{out} c^4}{\kappa_{out}^4 + c^4}}_{j_{out}} \quad [4]$$

$$\frac{dc_{tot}}{dt} = \underbrace{j_0 + j_a a}_{j_{in}} - \underbrace{\frac{v_{out} c^4}{\kappa_{out}^4 + c^4}}_{j_{out}}, \quad [5]$$

where $j_{rel}$ and $j_{serca}$ are endoplasmic reticulum $Ca^{2+}$ fluxes, and $j_{in}$ and $j_{out}$ are plasma membrane $Ca^{2+}$ fluxes. The functions $a_\infty$, $s_\infty$, $\theta_\infty$ and $\tau_\theta$ that appear in the epnea subsystem (Eqs. 1–3) are

$$a_\infty(x, c) = \frac{1}{1 + e^{4(\gamma_a - x)/k_a}} + \frac{\lambda_c}{1 + e^{4(\gamma_c - c)/k_c}} \quad [6]$$

$$s_\infty(x) = \frac{1}{1 + e^{4(\gamma_s - x)/k_s}} \quad [7]$$

$$\theta_\infty(x) = \frac{1}{1 + e^{4(\gamma_\theta - x)/k_\theta}} \quad [8]$$

$$\tau_\theta(x) = \frac{\tau_\theta^{max} - \tau_\theta^{min}}{1 + e^{4(\gamma_{\tau_\theta} - x)/k_{\tau_\theta}}} + \tau_\theta^{min}. \quad [9]$$

The first state variable, denoted by $a$, is the network activity of the preBötC. This dimensionless quantity takes values between 0 (no activity) and $1 + \lambda_c$ (maximum population firing rate). The dimensionless variables, $s$ and $\theta$, model the dynamics of synaptic depression and cellular adaptation, respectively. The variable $c$ is the cytosolic free $Ca^{2+}$ concentration ($c = [Ca^{2+}]$). The variable $c_{tot}$ is the total $[Ca^{2+}]$, a quantity dominated by intracellular stores such as the endoplasmic reticulum (ER). Writing ER $[Ca^{2+}]$ as $c_{er}$, the total $[Ca^{2+}]$ is defined as $c_{tot} = c + \rho c_{er}$, where where $\rho$ is an ER-to-cytosol volume ratio that accounts for the $Ca^{2+}$ buffering capacity of both compartments. There are two algebraic functions that occur in Eq. 4. The first expresses the ER $[Ca^{2+}]$ in terms of the cytosolic and total $[Ca^{2+}]$,

$$c_{er} = \frac{c_{tot} - c}{\rho}. \quad [10]$$

The second function is the bell-shaped open probability of the $IP_3$ receptor ($IP_3R$) as a function of cytosolic $[Ca^{2+}]$,

$$f_{open}(c) = \frac{1}{1 + e^{4(\gamma_m - c)/k_m}} \cdot \frac{1}{1 + e^{4(\gamma_h - c)/k_h}}. \quad [11]$$

See Appendix Tables 1–2 for a description of parameters and their standard values.

**Overview of SI Appendix:** The remainder of this supplemental text presents the various components of the mathematical model of the inspiratory and sigh rhythms (Eqs. 1–11). Section 1 focuses on the activity model of episodic inspiratory burst generation. Section 2 presents the dynamical model of slow $Ca^{2+}$ oscillations that drive the sigh rhythm. Section 3 gives details regarding the coupling and interaction of these two subsystems and discusses how this mathematical model of the inspiratory-sigh rhythm informed the experimental work discussed in the main text. Section 4 describes stochastic aspects of the inspiratory model. Section 5 compares the activity model of inspiratory rhythmogenic preBötzinger complex neurons with a spiking network model that includes $Ca^{2+}$ dynamics for $N$ distinct neurons with excitatory interactions mediated by a physiologically plausible network topology.

#### 1. Activity model of episodic bursting

Episodic bursting of the preBötC may be modeled as a two-variable dynamical system,

$$\tau_a \frac{da}{dt} = a_\infty(w \cdot s \cdot a) - a \qquad [12]$$

$$\tau_s \frac{ds}{dt} = s_\infty(a) - s\,, \qquad [13]$$

where the state variable $a$ is the network activity and $s$ accounts for the dynamics of synaptic depression ($s = 1$ indicates the absence of depression, while $s = 0$ corresponds to full depression). In Eqs. 12–13, the

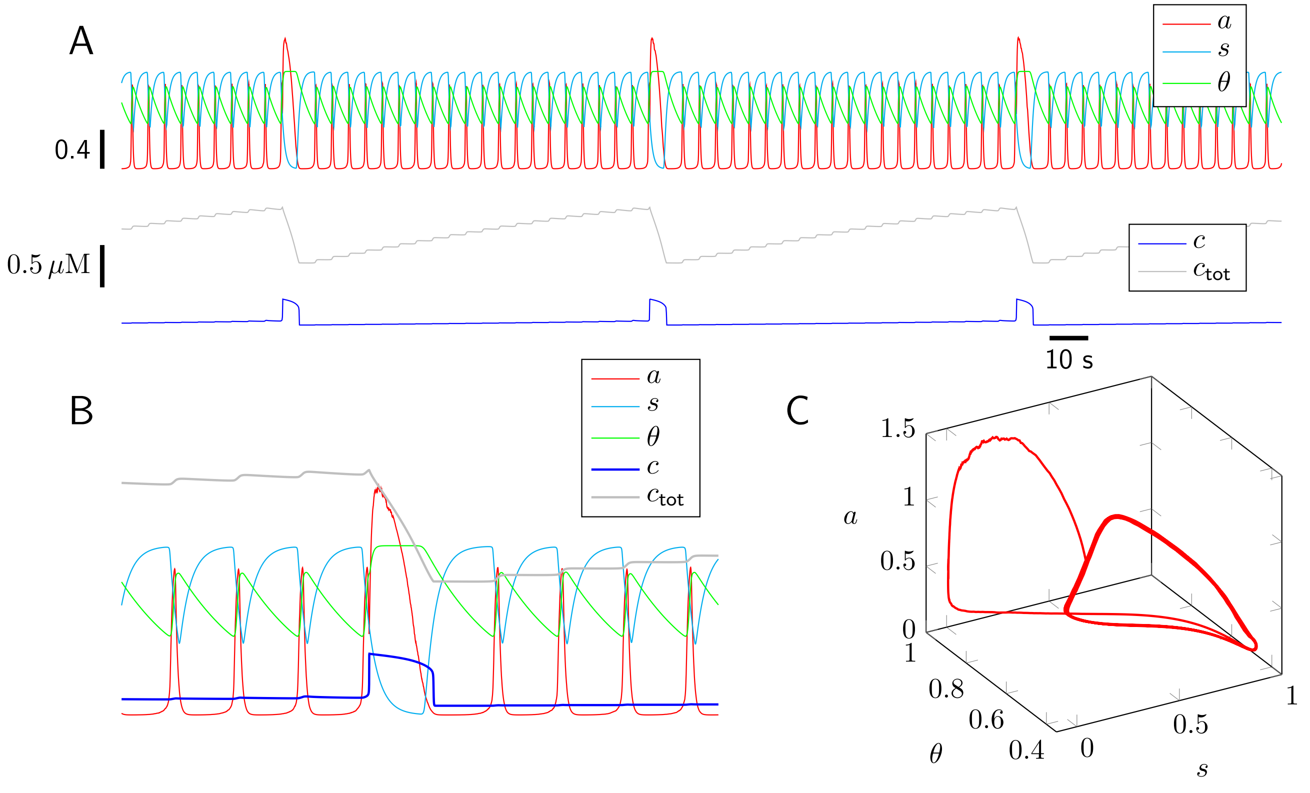


**Appendix Figure 1.** Representative simulation of the coupled inspiratory and sigh rhythms. (A) Dynamics of the five dependent variables of the ODE model. (B) Expanded view that includes a single sigh event that emerges shortly after the onset of inspiratory burst. (C) Three dimensional phase space for the ($a$, $s$, $\theta$) subsystem. Parameters as in Appendix Tables 1–2.

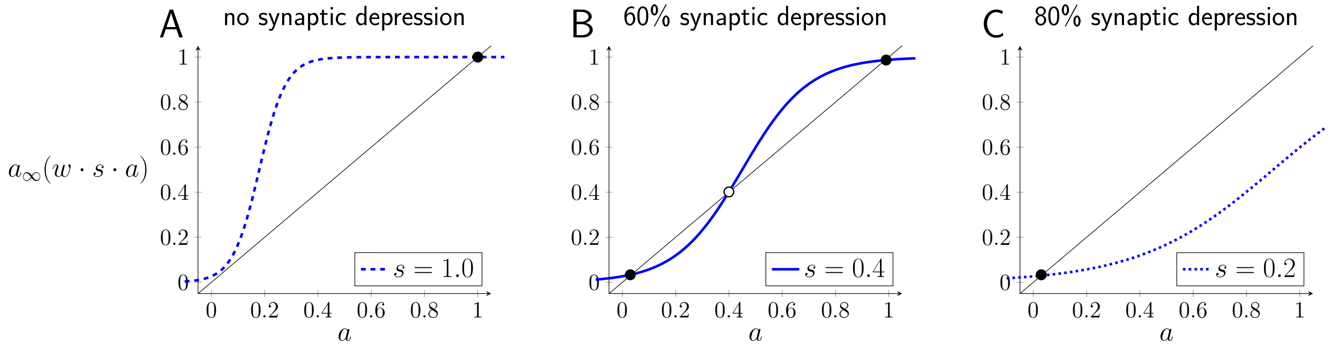


**Appendix Figure 2.** The recurrent excitatory network has steady-state population firing rates given by solutions of Eq. 16. The number of steady states depends on synaptic depression ($s$), which takes values between 0 (complete depression) and 1 (no depression). Stable and unstable steady states are shown by filled and open circles, respectively. (A) In the absence of synaptic depression there is a steady state near the maximal population firing rate ($a \approx 1$) at the intersection of $a_\infty$ (solid blue curve) and the 45° line (black) for $a = a_\infty(wsa)$. (B) For intermediate synaptic depression the network is bistable. (C) For highly depressed synapses there is a stable steady state at low population firing rate ($a \approx 0$). Parameters: $w = 1$, $\gamma_a = 0.18$, $k_a = 0.2$.

functions $a_\infty$ and $s_\infty$ are increasing sigmoids,

$$a_\infty(x) = \frac{1}{1 + e^{4(\gamma_a - x)/k_a}} \qquad [14]$$

$$s_\infty(x) = \frac{1}{1 + e^{4(\gamma_s - x)/k_s}} . \qquad [15]$$

Because $k_a$ is positive, $a_\infty$ is a monotone increasing function of $x$, which in Eq. 12 is the aggregate synaptic drive, calculated as the triple product $w \cdot s \cdot a$ (the standard value for the synaptic gain is $w = 1$, see Appendix Table 1). Conversely, $k_s$ is negative; hence, $s_\infty$ in Eq. 13 is monotone decreasing function of the network activity ($a$). The 4 that appears in Eqs. 14 and 15 makes $k_a$ and $k_s$ the inverse of the slope of $a_\infty$ and $s_\infty$ at their respective half-maxima. At steady state, the network activity ($a$) solves

$$\bar{a} = a_\infty(w \cdot \bar{s} \cdot \bar{a}) . \qquad [16]$$

Using the default value for the gain of recurrent excitation ($w = 1$), Appendix Fig. 2 shows that the network activity can have 1–3 steady states (solutions of Eq. 16) depending on the amount of synaptic depression ($s$). For intermediate values of $s$, the excitatory network is bistable (Appendix Fig. 2B).

**Dynamics of synaptic depression.** Appendix Fig. 3 shows representative phase planes that illustrate how the two-variable model (Eqs. 12–13) exhibits episodic network activity. The blue curve shows the $s$ nullcline, i.e., the loci (curves) in the $(s, a)$-plane for which $ds/dt = 0$,

$$s \text{ nullcline:} \quad s = s_\infty(a) , \qquad [17]$$

with $s_\infty$ given by Eq. 15. The $a$ nullcline ($da/dt = 0$) is given by the implicit expression $a = a_\infty(w \cdot s \cdot a)$, which is equivalent to the explicit expression

$$a \text{ nullcline:} \quad s = \frac{4\gamma_a - k_a \ln\left[(1-a)/a\right]}{4wa} . \qquad [18]$$

Appendix Fig. 3 shows that the Tabak-Rinzel-like model of the preBötC may exhibit two distinct forms of excitability and/or oscillations (namely, type 1 and type 2) depending on the parameters used; for review see (1). Appendix Fig. 3A shows the nullclines that give rise to type-2 oscillations; while Appendix Fig. 3B shows nullclines associated with type-1 excitability. The bifurcation diagram of Appendix Fig. 3C show oscillations emerging via a Hopf bifurcation (type 2). The bifurcation diagram of Appendix Fig. 3D show oscillations that arise from a saddle-node on an invariant circle bifurcation (SNIC, type 1).

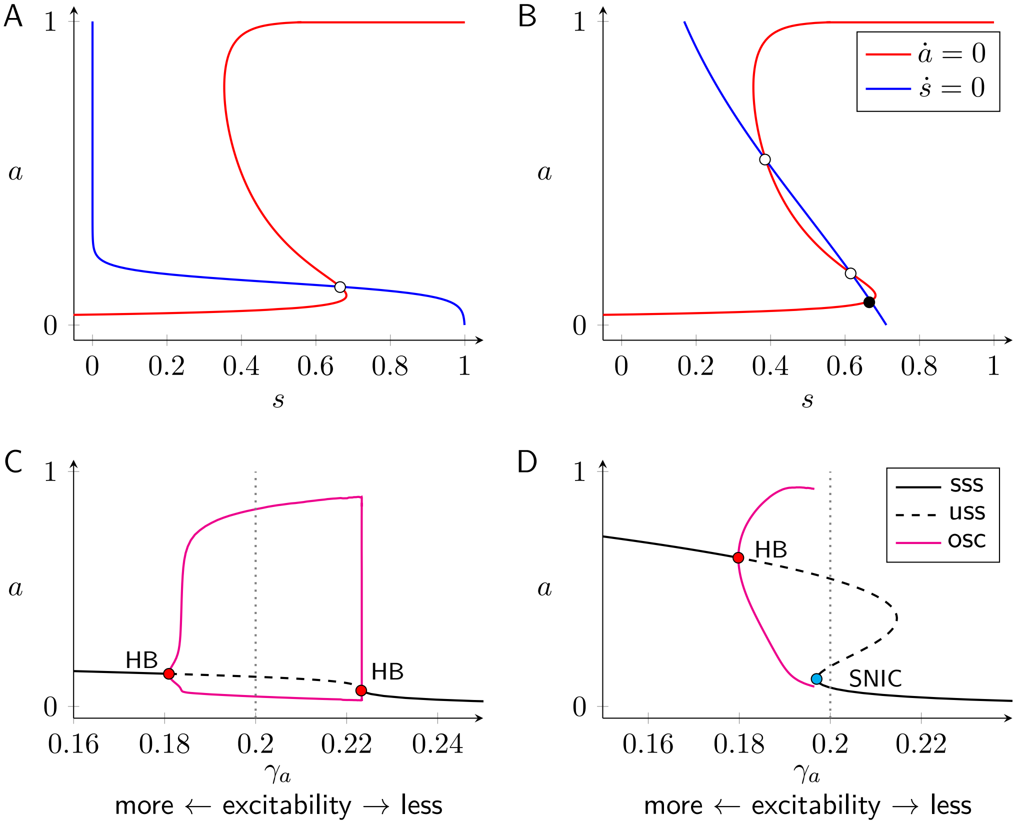


**Appendix Figure 3.** Nullclines and bifurcation diagrams for the two-variable model of inspiratory network activity (Eqs. 17–18). (A,B) Filled and open circles are stable and unstable steady states, respectively. (A) Nullclines for type-2 oscillations; $\gamma_s = 0.14$, $k_s = -0.08$. (B) Nullclines for type-1 excitability; $\gamma_s = 0.3$, $k_s = -1.0$. Other parameters: $w = 1$, $\tau_a = 10$ s, $\gamma_a = 0.2$, $k_a = 0.2$. (C,D) Bifurcation diagrams show emergence of oscillations as the average firing threshold ($\gamma_a$) is varied (lesser values of $\gamma_a$ lead to greater excitability, see Eq. 14). Filled circles show critical points: HB, Hopf bifurcation; SNIC, saddle-node on an invariant circle bifurcation. Stable steady states (sss) are shown solid black. Unstable steady states (uss) are shown dashed black. Minimum and maximum of limit cycle oscillations (osc) are shown magenta. Vertical dotted lines show value of $\gamma_a$ used in the corresponding phase diagrams (located above).

**Inspiratory dynamics with two variables (one fast and one slow).** How well does the two variable minimal model (Eqs. 12–15) of the inspiratory rhythm emulate *in vitro* preBötC recordings? Appendix Fig. 4A (left) shows a representative in vitro field recording of preBötC electrical activity. The scatter plot and histogram (right panel) illustrate two salient empirical characteristics of this rhythm. (1) The duration of the preceding interval has no influence on the size of a burst ($r^2 = 0.01$). (2) The distribution of inter-burst intervals is bell-shaped ($\mu = 5.4\,\text{s}$, $\sigma = 2.0\,\text{s}$).

Appendix Fig. 4B shows trajectories of episodic activity ($a$) and synaptic depression ($s$) for the inspiratory model with parameters leading to type-2 excitability. For realism, additive Gaussian white noise, $\tilde{\xi}(t)$, has been introduced to Eq. 12, as follows:

$$\tau_a \frac{da}{dt} = a_\infty(w \cdot s \cdot a) - a + \tilde{\xi}(t) \quad [19]$$

$$\tau_s \frac{ds}{dt} = s_\infty(a) - s\,. \quad [20]$$

The noise term, $\tilde{\xi}(t)$, is a rapidly fluctuating function of time with mean zero, $\langle \tilde{\xi}(t) \rangle = 0$, and variance that depends on system state (see Section 4). In the type-2 parameter regime, the model exhibits stochastic oscillations with inter-burst intervals distributed in a bell-shaped manner that is consistent with experiment (Appendix Fig. 4B histogram, $\mu = 4.1\,\text{s}$, $\sigma = 0.21\,\text{s}$). However, this episodic activity exhibits an unrealistic

**Appendix Table 1. Standard parameters for inspiratory model (dimensionless).**

| Symbol | Definition | Value |
|---|---|---|
| $w$ | network connectivity | $1$ |
| $\gamma_a$ | synaptic drive for half-maximal network activity $a_\infty$ (i.e., average firing threshold) | $-0.3$ |
| $k_a$ | reciprocal of slope of $a_\infty$ at half maximum | $0.2$ |
| $\tau_a$ | network recruitment time constant | $0.15$ |
| $\gamma_s$ | network activity for half-maximal $s_\infty$ | $0.14$ |
| $k_s$ | reciprocal of slope of $s_\infty$ at half maximum | $-0.08$ |
| $\tau_s$ | time constant of synaptic depression | $0.75$ |
| $\gamma_\theta$ | network activity for half-maximal $\theta_\infty$ | $0.15$ |
| $k_\theta$ | reciprocal of slope of $\theta_\infty$ at half maximum | $0.2$ |
| $\tau_\theta^{max}$ | maximum of time constant of cellular adaptation | $6$ |
| $\tau_\theta^{min}$ | minimum of time constant of cellular adaptation | $0.15$ |
| $\gamma_{\tau_\theta}$ | network activity for half-maximal $\tau_\theta(a)$ | $0.3$ |
| $k_{\tau_\theta}$ | reciprocal of slope of $\tau_\theta(a)$ at half maximum | $-0.5$ |

positive correlation between preceding interval and burst size (scatter plot, $r^2 = 0.21$). This correlation is due to fluctuations in network activity that are visible at the bottom knee of the $a$ nullcline (Appendix Fig. 4, phase plane). Variation in the level of synaptic depression when bursts are initiated leads to correlation between burst size and the duration of the preceding inter-burst interval (2).

For comparison, Appendix Fig. 4C shows simulations of the inspiratory model with parameters leading to stochastic type-1 excitability. In this parameter regime, the system is subcritical to a SNIC bifurcation. Because synaptic depression recovers to the same value before the onset of each network burst, the burst amplitudes are uncorrelated with the duration of preceding inter-event intervals ($r^2 < 0.01$). Bursts do not occur unless triggered by sufficiently large fluctuations (rare events); hence, the inter-burst intervals are distributed in an exponential fashion ($\mu = 6.8\,\mathrm{s}$), albeit shifted by the time required for recovery of synaptic depression. Because this inter-burst interval distribution is inconsistent with in vitro experimental observations (Fig. 4A of main text), type-1 stochastic excitability is an unrealistic starting point for our activity model of episodic bursting in the preBötC.

In summary, the simulations of Appendix Fig. 4 show that the two-variable model of inspiratory dynamics is unrealistic in the following sense. If parameters are chosen for stochastic type-2 oscillations, the inter-burst intervals are distributed in a realistic manner, but the positive correlation between the burst amplitude and the duration of the preceding interval is inconsistent with experiment. Conversely, if parameters are chosen for type-1 stochastic excitability, there is no correlation between burst amplitude and the duration of the preceding interval (consistent with experiment). But in this case the distribution of the inter-event intervals exhibited by the model is unrealistic (more variable than observed in experiment).

**Inspiratory dynamics with three variables (one fast and two slow).** The model development and analysis of the previous section suggests that the mechanism(s) that determines the inter-burst interval for inspiratory rhythm is distinct from the synaptic depression mechanism that is responsible for burst termination. For this reason, our activity model of the inspiratory subsystem includes a second slow variable, $\theta$, representing activity-dependent cellular adaptation. The three-variable model is

$$\tau_a \frac{da}{dt} = a_\infty(w \cdot s \cdot a - \theta) - a + \tilde{\xi}(t) \qquad [21]$$

$$\tau_s \frac{ds}{dt} = s_\infty(a) - s \qquad [22]$$

$$\tau_\theta(a) \frac{d\theta}{dt} = \theta_\infty(a) - \theta\,, \qquad [23]$$

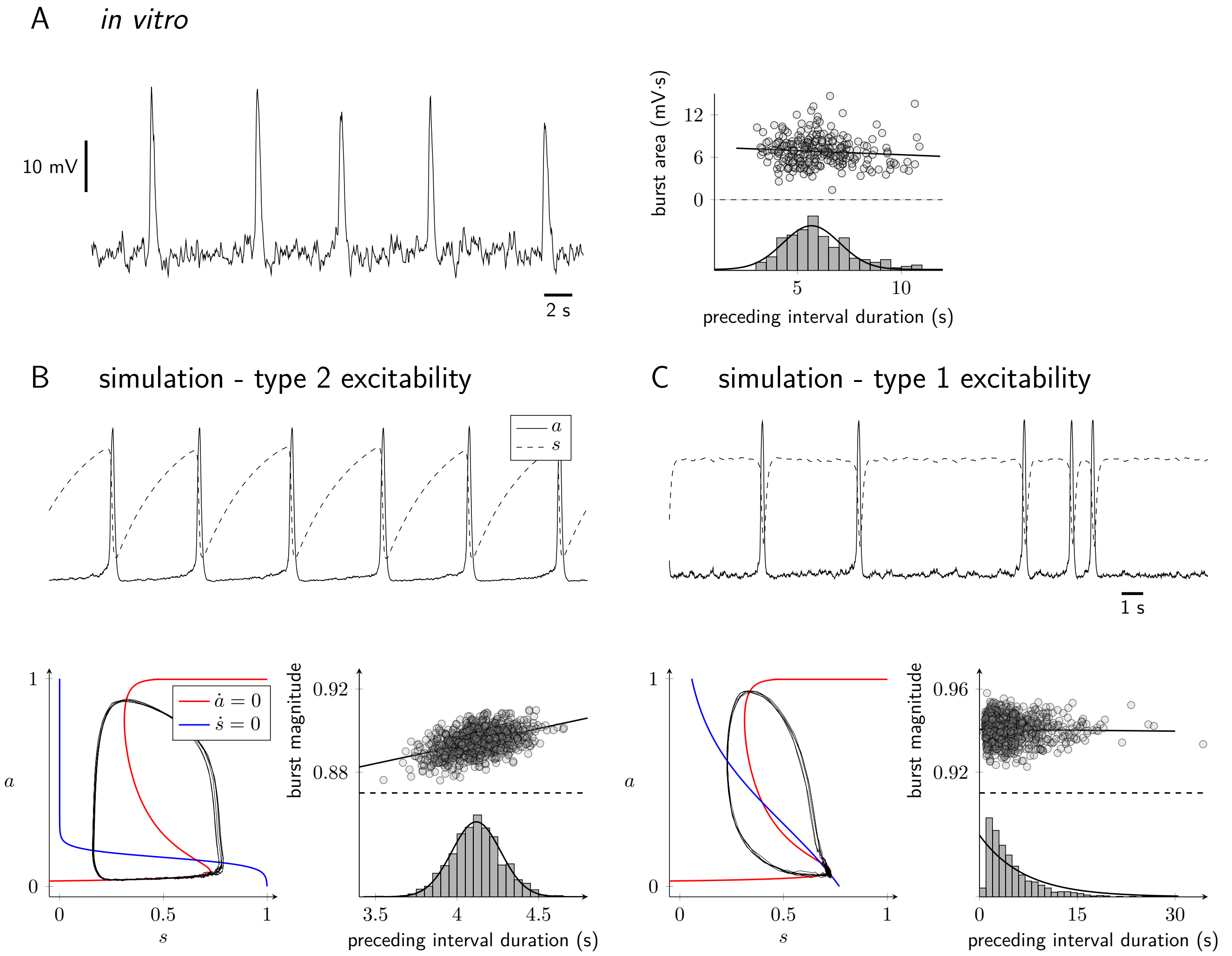


**Appendix Figure 4.** Limitations of the two-variable inspiratory model. (A) Field recording of preBötC activity. Right: Scatter plot of the relationship between burst size (area) and the duration of the preceding inter-burst interval. The histogram shows the distribution of inter-event intervals. (B) Simulations using the two-variable model with parameters giving type-2 stochastic oscillations. (C) Simulations using the two variable model with parameters giving type-1 stochastic excitability. Parameters are as in Appendix Table 1 except for type-2 excitability: $\gamma_a = 0.18$, $\tau_a = 0.15$ s; and type-1 excitability: $\gamma_a = 0.18$, $\tau_a = 0.03$ s, $k_s = -1$, $\gamma_s = 0.3$, $\tau_s = 0.1$ s.

with $a_\infty$ and $s_\infty$ given by Eqs. 14–15. The steady-state level of cellular adaptation,

$$\theta_\infty(a) = \frac{1}{1 + e^{4(\gamma_\theta - a)/k_\theta}}\,, \qquad [24]$$

is an increasing function of network activity ($k_\theta > 0$). The time constant for cellular adaptation, $\tau_\theta$, is also a function of network activity,

$$\tau_\theta(a) = \frac{\tau_\theta^{max} - \tau_\theta^{min}}{1 + e^{4(\gamma_{\tau_\theta} - a)/k_{\tau_\theta}}} + \tau_\theta^{min}\,. \qquad [25]$$

Parameters are chosen to ensure that cellular adaptation accumulates rapidly in the active phase (large $a$), but recovers slowly during the silent phase when network activity is low (Appendix Table 1). Appendix Fig. 5A shows a representative simulation of the three-variable inspiratory model exhibiting episodic bursting. In the three-variable model, synaptic depression terminates the bursts, but burst onset is determined by the recovery of cellular adaptation ($\theta$ must be sufficiently small), consistent with empirical data (3–5). In particular, (1) the inter-burst interval distribution is normally distributed with realistic mean and variance

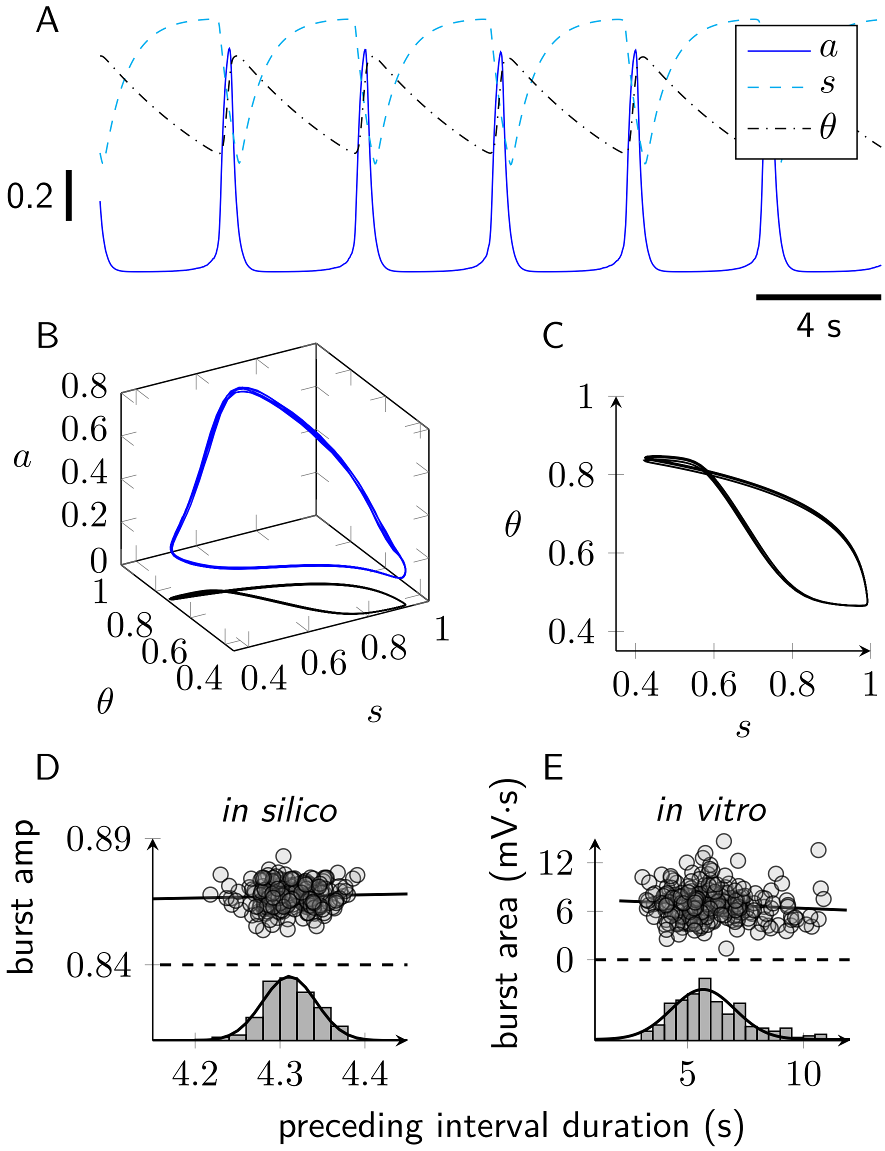


**Appendix Figure 5.** Three-variable model of episodic network activity gives realistic inspiratory dynamics (see Eqs. 21–23). (A) Representative trajectory of the three-state model showing network activity ($a$) and the dynamics of two slow variables: synaptic depression ($s$) and cellular adaptation ($\theta$). (B) Trajectory in 3d phase space (blue curve, left) with projection emphasizing the relationship between synaptic depression and cellular adaption ($s$ and $\theta$, black curve, right). (C) Comparison of burst amplitude and inter-burst interval duration according to the model (in silico) and experiment (in vitro), reproduced from Appendix Fig. 4A).

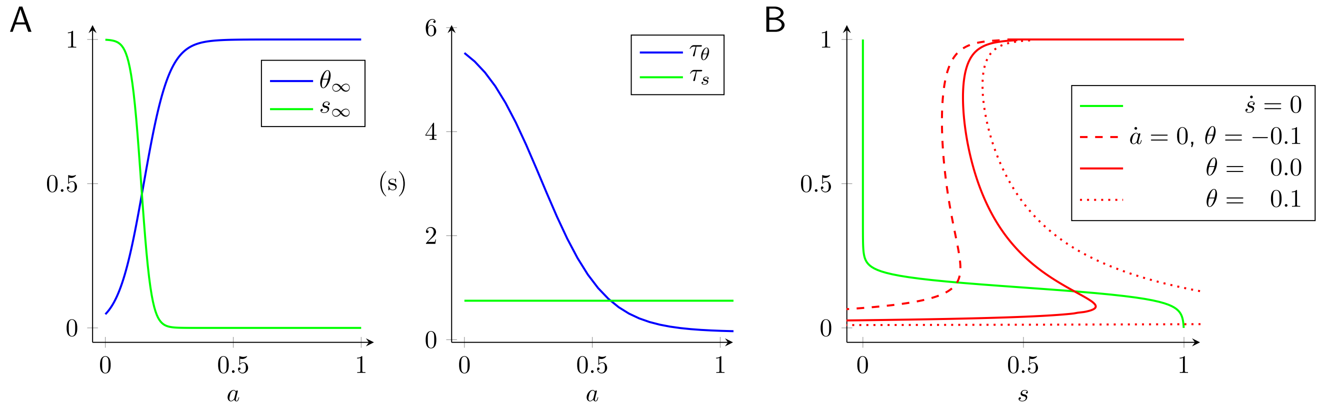


**Appendix Figure 6.** (A) Dynamics of synaptic depression ($s$) and cellular adaption ($\theta$) depend upon network activity ($a$) via steady-state functions, $s_\infty$ and $\theta_\infty$, and time constants, $\tau_s$ and $\tau_\theta$. (B) Steady-state network activity for three different values of the variable representing cellular adaptation ($\theta$, see legend).

($\mu = 4.3\,\text{s}$, $\sigma = 0.044\,\text{s}$), and (2) there is no correlation between burst amplitude and the duration of the preceding inter-burst interval ($r^2 < 0.01$) (cf. Appendix Fig. 5D and E).

#### 2. $Ca^{2+}$ handling and the sigh rhythm

Appendix Fig. 1 shows a representative simulation of the coupled inspiratory and sigh rhythms with three sigh events. These occur because the inspiratory subsystem ($a$, $s$, $\theta$) is coupled to oscillatory dynamics for intracellular $Ca^{2+}$ ($c$, $c_{tot}$) that periodically evoke a $Ca^{2+}$-dependent increase in network activity (Eq. 6). The $Ca^{2+}$ subsystem is the following two ODEs,

$$\frac{dc}{dt} = \underbrace{[v_{ip3r} f_{open}(c) + v_{leak}]\,[c_{er}(c, c_{tot}) - c]}_{j_{rel}} - \underbrace{\frac{v_{serca} c^2}{\kappa_{serca}^2 + c^2}}_{j_{serca}} + \underbrace{j_0 + j_a a}_{j_{in}} - \underbrace{\frac{v_{out} c^4}{\kappa_{out}^4 + c^4}}_{j_{out}} \quad [26]$$

$$\frac{dc_{tot}}{dt} = \underbrace{j_0 + j_a a}_{j_{in}} - \underbrace{\frac{v_{out} c^4}{\kappa_{out}^4 + c^4}}_{j_{out}}, \quad [27]$$

where the variable $c$ is the cytosolic [$Ca^{2+}$], and $c_{tot}$ is the total intracellular [$Ca^{2+}$] that includes contributions from the cytosol but is dominated by intracellular stores such as the endoplasmic reticulum (ER). See (6, 7) for review of $Ca^{2+}$ dynamics modeled in this fashion.

The parameters $v_{ip3r}$ and $v_{leak}$ in Eq. 26 are rate constants for $Ca^{2+}$-induced $Ca^{2+}$ release and a passive leak, both with a driving force given by the concentration gradient across the ER membrane ($c_{er} - c$). The parameter $v_{serca}$ is the maximal activity of a sarco-endoplasmic reticulum $Ca^{2+}$ ATPase (SERCA) reuptake flux given by a sigmoidal Hill expression with dissociation constant $\kappa_{serca}$. The parameter $v_{out}$ is the maximal activity of the sigmoidal expression $j_{out}(c)$ representing extrusion of $Ca^{2+}$ by plasma membrane $Ca^{2+}$ ATPases (PMCA). The terms $j_0 + j_a a$ in Eqs. 26–27 model $Ca^{2+}$ influx as a linear function of the network activity $a$ where $j_a$ is a propotionality constant and $j_0$ is the background rate. The algebraic functions that occur in Eqs. 26 and 27) are given by $c_{er} = (c_{tot} - c)/\rho$ and Eq. 11.

**Bistability in a closed cell model of $Ca^{2+}$ handling.** The relaxation oscillator dynamics of the $Ca^{2+}$ subsystem that drive the sigh rhythm can be understood by considering how the dynamics of cytosolic [$Ca^{2+}$] depends on total cell $Ca^{2+}$ ($c_{tot}$), which is the slow variable in Eqs. 26–27. If this slow variable were

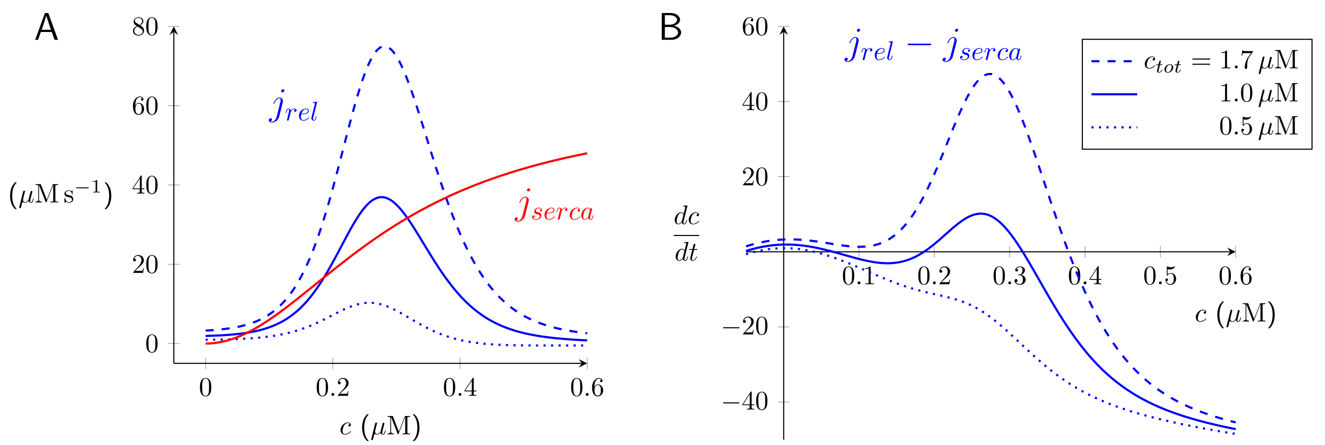


**Appendix Figure 7.** In the closed cell model of $Ca^{2+}$ handling (Eq. 28) there are no plasma membrane fluxes; consequently, the total $[Ca^{2+}]$ ($c_{tot}$) is parameter (not a state variable). (A) Fluxes associated to $IP_3$R-mediated $Ca^{2+}$ release ($j_{rel}$) and SERCA pumps ($j_{serca}$) as a function of cytosolic $Ca^{2+}$ concentration ($c$). The release flux $j_{rel}$ is an increasing function of $c_{tot}$ because it is proportional to the concentration gradient ($c_{er} - c$) where $c_{er} = (c_{tot} - c)/\rho$ (compare dotted, dashed, and solid curves; $c_{tot}$ as in legend in B). The reuptake flux $j_{serca}$ (shown red) is a function of $c$ but not $c_{tot}$. (B) Phase diagram of closed cell model of $Ca^{2+}$ handling (Eq. 28). Because plasma membrane fluxes are not included in the close cell model, $c_{tot}$ is constant (legend shows three values). Parameters as in Appendix Table 2.

constant, the following ODE for the closed cell model would apply,

$$\frac{dc}{dt} = h(c) \quad = \quad \underbrace{\left[v_{ip3r} f_{open}(c) + v_{leak}\right] (c_{er}(c) - c)}_{j_{rel}} - \underbrace{\frac{v_{serca} c^2}{\kappa_{serca}^2 + c^2}}_{j_{serca}} \quad \text{where} \quad c_{er} = \frac{c_{tot} - c}{\rho}. \qquad [28]$$

Using three different values for $c_{tot}$, Appendix Fig. 7 plots the net ER flux $h(c) = j_{rel} - j_{serca}$. For intermediate values of $c_{tot}$ (solid blue curve), $h(c)$ intersects the horizontal axis three times. Noting the slope $h'(c)$ evaluated at these three steady states, it is evident that the low and high steady states are stable while the intermediate steady state is unstable.

**Appendix Table 2. Standard parameters for $Ca^{2+}$ subsystem.**

| Symbol | Definition | Value | Units |
|---|---|---|---|
| $v_{ip3r}$ | rate constant of $Ca^{2+}$ release | 20 | $s^{-1}$ |
| $v_{leak}$ | rate constant of $Ca^{2+}$ leak | 0.25 | $s^{-1}$ |
| $v_{serca}$ | maximum rate of SERCA pumps | 60 | $\mu M\, s^{-1}$ |
| $\kappa_{serca}$ | half maximum for SERCA pumps | 0.3 | $\mu M$ |
| $\rho$ | ER/cytosol effective volume ratio | 0.15 | - |
| $\gamma_m$ | activation of intracellular $Ca^{2+}$ channels | 0.25 | $\mu M$ |
| $k_m$ | reciprocal of slope of $m_\infty$ at half maximum | 0.16 | $\mu M^{-1}$ |
| $\gamma_h$ | activation of intracellular $Ca^{2+}$ channels | 0.3 | $\mu M$ |
| $k_h$ | reciprocal of slope of $h_\infty$ at half maximum | $-0.24$ | $\mu M^{-1}$ |
| $j_0$ | constant $Ca^{2+}$ influx rate | 0 | $\mu M\, s^{-1}$ |
| $j_a$ | $Ca^{2+}$ influx rate proportionality constant | 0.1 | $\mu M\, s^{-1}$ |
| $v_{out}$ | maximum rate of PMCA pumps | 0.4 | $\mu M\, s^{-1}$ |
| $\kappa_{out}$ | half maximum for PMCA pumps | 0.3 | $\mu M$ |
| $\lambda_c$ | maximum $Ca^{2+}$-dependent increase of activity | 1.5 | - |
| $\gamma_c$ | threshold for $Ca^{2+}$-dependent increase of activity | 0.33 | $\mu M$ |
| $k_c$ | reciprocal slope of $Ca^{2+}$-dependent increase of activity | 0.05 | $\mu M$ |

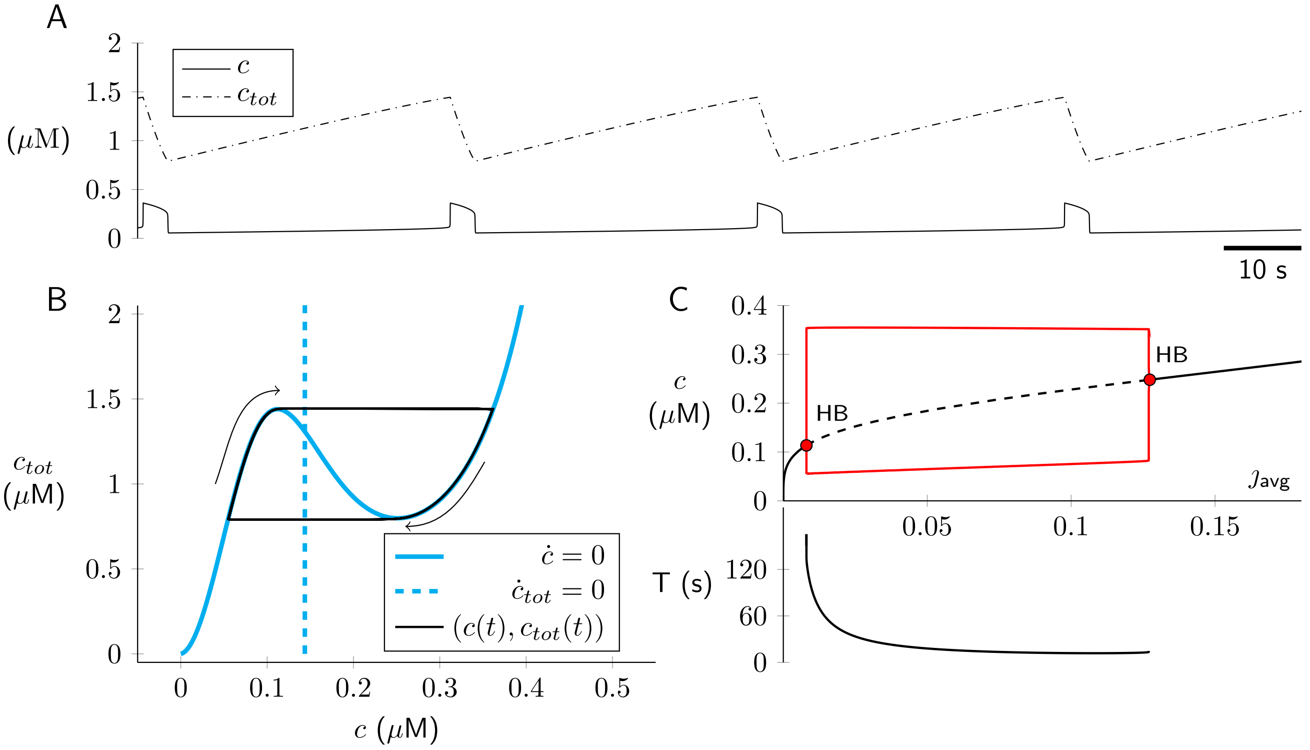


**Appendix Figure 8.** Open cell $Ca^{2+}$ dynamics given by solution of Eqs. 30–31. Parameters: $j_{in}^{avg} = 0.02\ \mu M\,s^{-1}$ and as in Appendix Table 2. (A) Type-2 relaxation oscillations ($c$ fast, $c_{tot}$ slow). (B) Phase plane, with $c$ and $c_{tot}$ nullclines (cyan solid and broken lines, respectively) and periodic solution (black). (C) Bifurcation diagram with $j_{in}^{avg}$ (average $Ca^{2+}$ influx rate) as bifurcation parameter. $T$ is period of relaxation oscillation.

**Relaxation oscillations in open cell model of $Ca^{2+}$ handling.** In the full model of coupled inspiratory and sigh rhythms (Eqs. 1–11), the fluxes representing $Ca^{2+}$ release and reuptake from the ER (Eq. 28) are augmented by plasma membrane fluxes, $j_{pm} = j_{in} - j_{out}$ where $j_{in} = j_0 + j_a a$ and $j_{out} = v_{out}c^4/(\kappa_{out}^4 + c^4)$. By averaging the $Ca^{2+}$ influx over cycles of episodic bursting with period $T$, one may calculate an effective $Ca^{2+}$ influx rate, given by

$$j_{in}^{avg} = \frac{1}{T}\int_0^T [j_0 + j_a a(t)]\, dt\,, \qquad [29]$$

that no longer depends on the network activity ($a$). The resulting open cell model is

$$\frac{dc}{dt} = \underbrace{[v_{ip3r} f_{open}(c) + v_{leak}]\,[c_{er} - c]}_{j_{rel}} - \underbrace{\frac{v_{serca}c^2}{\kappa_{serca}^2 + c^2}}_{j_{serca}} + j_{in}^{avg} - \underbrace{\frac{v_{out}c^4}{\kappa_{out}^4 + c^4}}_{j_{out}} \qquad [30]$$

$$\frac{dc_{tot}}{dt} = j_{in}^{avg} - \underbrace{\frac{v_{out}c^4}{\kappa_{out}^4 + c^4}}_{j_{out}}\,. \qquad [31]$$

Appendix Fig. 8 (top) shows the relaxation oscillator dynamics of this open cell model using standard parameters (Appendix Table 2) and $j_{in}^{avg} = 0.02\,\mu M\,s^{-1}$. The oscillation period is on the order of minutes and the system spends most of its time in the *down* state with low cytosolic $Ca^{2+}$ (0.05–0.11 $\mu$M) and slowly increasing total cell $Ca^{2+}$, $c_{tot}$, which implies slowly increasing ER $Ca^{2+}$, because $c_{er} = (c_{tot} - c)/\rho$. The phase plane of Appendix Fig. 8 (bottom left) shows the $c$ and $c_{tot}$ nullclines that are found by setting the left sides of Eqs. 30–31 to zero. The $c$ nullcline is $N$-shaped and has two extrema (knees). The $c_{tot}$

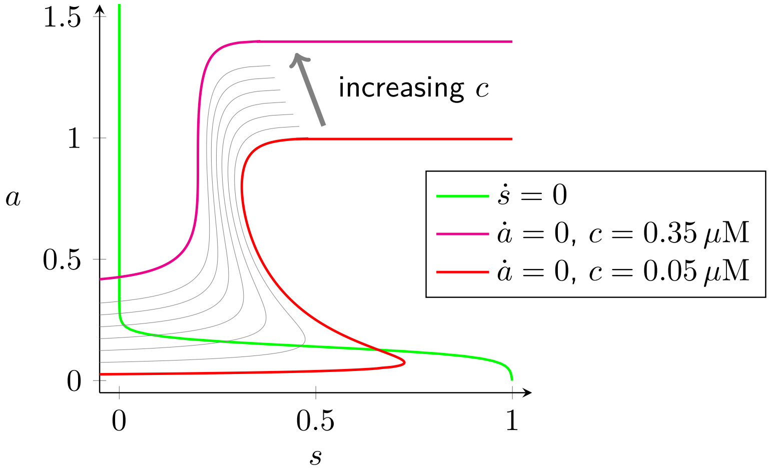


**Appendix Figure 9.** The influence of cytosolic $[\mathrm{Ca}^{2+}]$ ($c$, gray arrow) on the network activity nullcline ($\dot{a} = 0$) given by $a = a_\infty(wsa - \theta, c)$ (Eq. 33). For fixed $\theta$ and $c$ (legend), the nullcline $a = a_\infty(s)$ is given by Eq. 34. Parameters: $w = 1$, $\theta = 0.18 - \gamma_a = 0.48$, $\lambda_c = 0.8$, $\gamma_c = 0.35$, $k_c = 0.05$ and as in Appendix Tables 1–2. This gives $a^c_\infty(c) = 0$ to $0.4$ for $c = 0.05$ to $0.35\,\mu$M.

nullcline is a vertical line located at the value of $c$ which solves $j_{in}^{avg} = j_{out}(c) = v_{out}c^4/(\kappa_{out}^4 + c^4)$. Note how the separation of time scales for $c$ and $c_{tot}$ leads to a periodic solution (black trajectory) that tracks the lower or upper branch of the $c$ nullcline except for two brief excursions between branches when the trajectory passes over a knee of the $c$ nullcline. Appendix Fig. 8 (bottom right) shows a bifurcation diagram for Eqs. 30–31. Type-2 relaxation oscillations that originate via Hopf bifurcations are observed for a wide range of average $\mathrm{Ca}^{2+}$ influx rates ($j_{in}^{avg}$). Given the separation of time scales, oscillations occur at values of $j_{in}^{avg}$ that cause the $c$ nullcline to intersect the $c_{tot}$ nullcline between the knees.

#### 3. Coupling of the inspiratory and sigh rhythms

In the model of inspiratory and sigh rhythmogenesis (Eqs. 1–11), the fast ($a$, $s$, $\theta$) and slow ($c$, $c_{tot}$) subsystems are bi-directionally coupled in a manner that creates the inspiratory/sigh dynamics (Appendix Fig. 1A). Fast to slow: Episodic network activity ($a$) influences dynamics of intracellular $\mathrm{Ca}^{2+}$ via the plasma membrane influx rate $j_{in} = j_0 + j_a a$ (Eqs. 4–5). Slow to fast: Cytosolic $\mathrm{Ca}^{2+}$ activates a cationic current (8) whose influence is modeled abstractly using a two-term network activity function,

$$a_\infty(x, c) = \underbrace{\frac{1}{1 + e^{4(\gamma_a - x)/k_a}}}_{a^x_\infty(x)} + \underbrace{\frac{\lambda_c}{1 + e^{4(\gamma_c - c)/k_c}}}_{a^c_\infty(c)}, \qquad [32]$$

where $a^x_\infty$ and $a^c_\infty$ depend on synaptic drive ($x$) and cytosolic $\mathrm{Ca}^{2+}$ ($c$), respectively. Thus, in the model of inspiratory and sigh rhythmogenesis, the steady-state network activity function in Eq. 1 is

$$a_\infty(w \cdot s \cdot a - \theta, c) = \underbrace{\frac{1}{1 + e^{4(\gamma_a + \theta - wsa)/k_a}}}_{a^x_\infty(wsa - \theta)} + \underbrace{\frac{\lambda_c}{1 + e^{4(\gamma_c - c)/k_c}}}_{a^c_\infty(c)}. \qquad [33]$$

Appendix Fig. 9 plots the network activity nullcline ($\dot{a} = 0$),

$$a \text{ nullcline:} \quad s = \frac{4(\gamma_a + \theta) - k_a \ln\left(\dfrac{1}{a - a^c_\infty(c)} - 1\right)}{4wa}, \qquad [34]$$

for a range of cytosolic [$Ca^{2+}$] ($0.05 \le c \le 0.35$ $\mu$M). Upon ER $Ca^{2+}$ release and during the active phase of the $Ca^{2+}$ oscillation, an upward shift of the $a$ nullcline (magenta) accounts for the increase in network activity mediated by $Ca^{2+}$-activated cationic current.

#### 4. Stochastic inspiratory models

Appendix Fig. 4 shows simulation of episodic activity ($a$) and synaptic depression ($s$) for the two-variable inspiratory model with parameters leading to type-2 stochastic oscillations (panel B) and type-1 stochastic excitability (panel C). The Gaussian white noise term $\tilde{\xi}(t)$ of Eq. 19 models stochastic action potential firing. It is a rapidly varying function of time with mean zero, $\langle \tilde{\xi}(t) \rangle = 0$, and two-time covariance,

$$\langle \tilde{\xi}(t)\tilde{\xi}(t') \rangle = \nu(a,s)\delta(t-t') \,. \qquad [35]$$

The state-dependent variance, $\nu(a,s)$, is derived by considering the rates of the forward and reverse elementary processes (9) implied by Eq. 19,

$$\frac{da}{dt} = \frac{a_\infty(w \cdot s \cdot a) - a}{\tau_a} = \frac{a_\infty(s,a) - a}{\tau_a} \,.$$

where $\lim_{x \to -\infty} a_\infty(x) = 0$ and $\lim_{x \to \infty} a_\infty(x) = 1$ (Eq. 6). To understand the second equality, recall that $a_\infty(w \cdot s \cdot a) = a_\infty(a,s)$ is a function of both state variables, but $w$ is a parameter. Defining $\alpha$ and $\beta$ to simultaneously solve $\tau_a = 1/(\alpha + \beta)$ and $a_\infty = \alpha/(\alpha+\beta)$, we see that Eq. 19 is equivalent to

$$\frac{da}{dt} = \alpha(a,s) \cdot (1-a) - \beta(a,s) \cdot a \,,$$

where $\alpha = a_\infty/\tau_a$ and $\beta = (1-a_\infty)/\tau_a$ are functions of $a$ and $s$ through $a_\infty(a,s)$. This equation is consistent with elementary processes as diagrammed below:

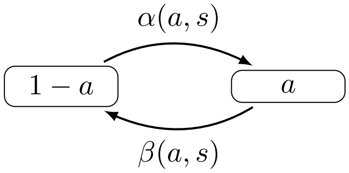


The variance used in Eq. 19 is the sum of the forward and reverse rates of transition, given by $\alpha(a,s) \cdot (1-a)$ and $\beta(a,s) \cdot a$, respectively. Thus,

$$\begin{aligned} \nu(a,s) &= \eta \left[ \alpha(a,s) \cdot (1-a) + \beta(a,s) \cdot a \right] \\ &= \frac{\eta}{\tau_a} \left[ a_\infty(a,s) \cdot (1-a) + (1 - a_\infty(a,s)) \cdot a \right] \,, \end{aligned}$$

where $\eta$ scales the strength of the noise. Larger networks with more neurons will have reduced noise compared to smaller networks.

#### 5. Spiking model of inspiratory and sigh rhythms

Sections 1–4 of this appendix have focused on the network activity model of inspiratory rhythmogenic preBötzinger complex neurons and the dependence of sigh breathing rhythm on intracellular $Ca^{2+}$ oscillations. As discussed briefly in the main text, the activity model approach can be compared and contrasted with models that include spiking and $Ca^{2+}$ dynamics for $N$ distinct neurons with excitatory interactions mediated by a physiologically plausible network topology. In Fig. S4, the voltage activity of each neuron is represented using the *theta model*, i.e., the Ermentrout-Kopell canonical model (10, 11),

$$\dot{\vartheta} = (1 - \cos\vartheta) + (1 + \cos\vartheta)\eta \qquad -\pi < \vartheta \le \pi \,, \tag{36}$$

where the state variable $\vartheta$ is an angle in radians (not to be confused with usage of $\theta$ in Section 1). The state space is circular (i.e., $-\pi$ and $\pi$ are identified) and the model produces a spike when $\vartheta$ increases across $\pi$. The parameter $\eta$ determines the intrinsic behavior of the model neuron (see Appendix Fig. 10). For $\eta < 0$ the neuron is excitable; there is a stable fixed point ($\vartheta^- < 0$, rest state) and unstable fixed point ($\vartheta^+ > 0$, threshold for action potential) where $\vartheta^\pm = \pm\cos^{-1}((1+\eta)/(1-\eta))$. For $\eta > 0$ the neuron spikes repetitively with period $T = \pi/\sqrt{\eta}$ and frequency $f = 1/T$.

Beginning with Eq. 36, a network model of inspiratory rhythmogenesis composed of $N$ spiking neurons may be constructed as follows:

$$\dot{\vartheta}_i = (1 - \cos\vartheta_i) + (1 + \cos\vartheta_i)\eta_i \quad \text{where} \quad \eta_i = \bar{\eta} + (\delta/\kappa_i^{in})\sum_j a_{ij}s_j(1 - n_j) + \tilde{\eta}_i(t) \tag{37}$$

$$\dot{s}_i = -s_i/\tau_s \tag{38}$$

$$\dot{n}_i = \alpha_n\, m_i\,(1 - n_i) - n_i/\tau_n \tag{39}$$

$$\dot{m}_i = -m_i/\tau_m \,. \tag{40}$$

In these equations, $\vartheta_i$ is the oscillatory phase of the $i$th neuron ($1 \le i \le N$). The excitatory drive for each neuron ($\eta_i$) is given by a sum over presynaptic neurons (Eq. 37). The state variables $s_i$, $n_i$ and $m_i$, which take values between 0 and 1, account for the the dynamics of excitatory synapses and synaptic depression. The factor $s_j(1 - n_j)$ in Eq. 37 is the synaptic activity of the $j$th neuron ($n_j = 0$ represents the absence of depression). When the $i$th neuron produces a spike, the values of $s_i$ and $m_i$ are both incremented as follows:

$$\vartheta_i(t) = \pi \implies \begin{cases} s_i(t^+) = s_i(t^-) + \alpha_s[1 - s_i(t^-)] \\ m_i(t^+) = m_i(t^-) + \alpha_m[1 - m_i(t^-)] \end{cases} . \tag{41}$$

Note that $n_i$ relaxes to 1 (complete depression) with exponential rate constant $\alpha_n m_i$. Because this rate is proportional to $m_i$, the dynamics of $n_i$ are second order (depression is not instantaneous).

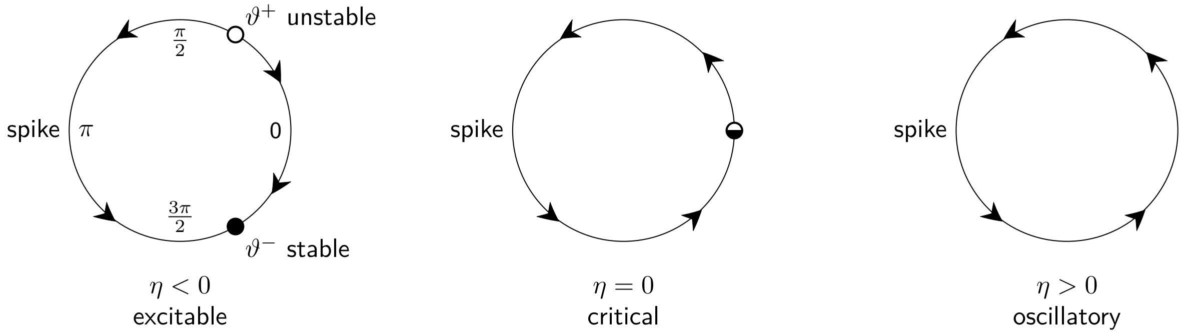


**Appendix Figure 10.** The Ermentrout-Kopell canonical model (Eq. 36) is excitable, critical, or oscillatory depending on the sign of the bifurcation parameter $\eta$.

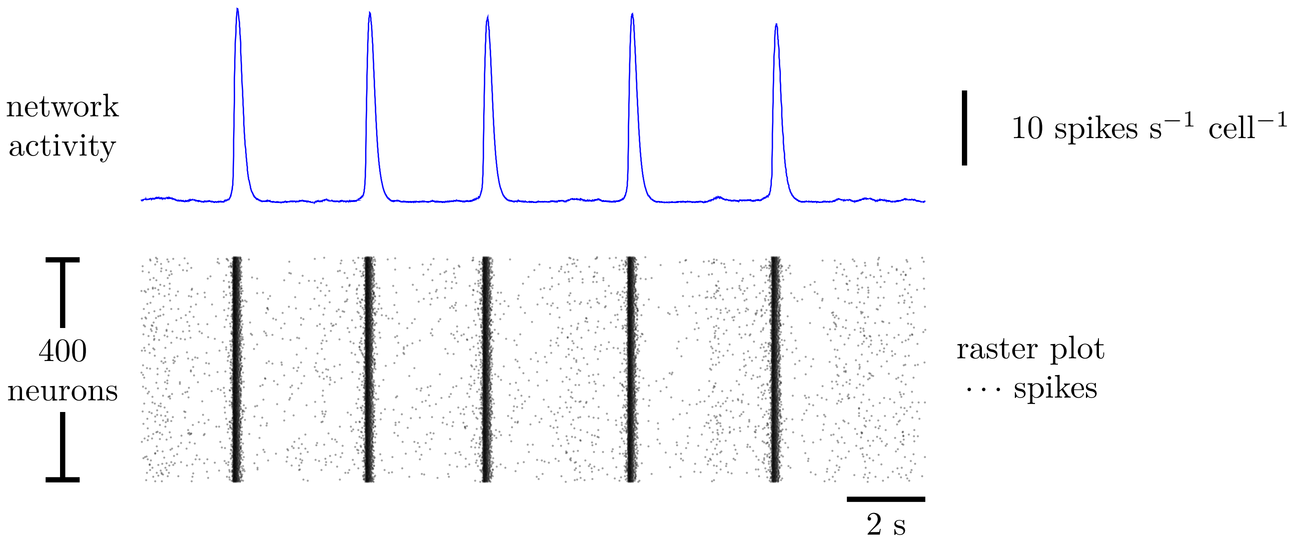


**Appendix Figure 11.** The spiking model of inspiratory rhythmogenesis (Eqs. 36–41). Erdös-Rényi-type network connectivity with $N = 400$ and $p = 0.065$ (see Section 5). Parameters as in first block of Appendix Table 3.

In Eq. 37, the network connectivity is given by an $N \times N$ adjacency matrix $A = (a_{ij})$. The element $a_{ij} > 0$ if neuron $i$ is postsynaptic to neuron $j$ and zero otherwise. Neurons are not self-excitatory ($a_{ii} = 0$). The in- and out-degrees of the $i$th neuron are given by $\kappa_i^{in} = \sum_j a_{ij}$ and $\kappa_i^{out} = \sum_j a_{ji}$, respectively. For a given simulation, the network connectivity is modeled as random directed Erdös-Réyni graph (12) with parameters $N = 400$ and $p = 0.065$. This value for $p$ greatly exceeds the threshold for connectedness given by $(\ln N)/N \approx 0.015$. The expected value for the in- and out-degree of each neuron is $\kappa = p(N-1) \approx 24$.

The excitatory drive for the $i$th neuron, $\eta_i = \bar{\eta} + (\delta/\kappa_i^{in}) \sum_j a_{ij} s_j (1 - n_j) + \tilde{\eta}_i(t)$, has three terms. The first term, $\bar{\eta}$ is negative so that the neurons intrinsically excitable (see Appendix Fig. 10). In the second term, the parameter $\delta$ sets the magnitude of recurrent excitation in the network. The factor of $1/\kappa_i^{in}$ scales the synaptic interactions so that every neuron receives the same amount of excitation when all of its presynaptic neurons are active. This ensures that the network dynamics will not depend on network size ($N$) provided it is sufficiently large. The third term is Gaussian white noise with mean zero $\langle \tilde{\eta}_i(t) \rangle = 0$ and two-time covariance $\langle \tilde{\eta}_i(t) \tilde{\eta}_i(t') \rangle = \nu \delta(t - t')$. Each neuron receives an independent stream of additive noise, that is, $\langle \tilde{\eta}_i \tilde{\eta}_j \rangle = 0$ for $i \neq j$.

Appendix Fig. 11 shows a representative simulation of the spiking model of inspiratory rhythmogenesis (Eqs. 36–41). To include the sigh rhythm, these equations are augmented with intracellular $Ca^{2+}$ dynamics for each neuron,

$$\frac{dc_i}{dt} = \underbrace{\left[v_{ip3r} f_{open}(c_i) + v_{leak}\right]\left[c_i^{er}(c_i, c_i^{tot}) - c_i\right]}_{j_i^{rel}} - \underbrace{\frac{v_{serca} c_i^2}{\kappa_{serca}^2 + c_i^2}}_{j_i^{serca}} + \underbrace{\jmath_0 + \jmath_\eta \eta_i}_{j_i^{in}} - \underbrace{\frac{v_{out} c_i^4}{\kappa_{out}^4 + c_i^4}}_{j_i^{out}} \quad [42]$$

$$\frac{dc_i^{tot}}{dt} = \underbrace{\jmath_0 + \jmath_\eta \eta_i}_{j_i^{in}} - \underbrace{\frac{v_{out} c_i^4}{\kappa_{out}^4 + c_i^4}}_{j_i^{out}}, \quad [43]$$

where $c_i^{er} = (c_i^{tot} - c_i)/\rho$. In these equations, $c_i$ and $c_i^{tot}$ denote the cytosolic and total $[Ca^{2+}]$ of the $i$th neuron (compare Eqs. 26–27). The $Ca^{2+}$ influx term, $j_i^{in} = \jmath_0 + \jmath_\eta \eta_i$, that appears in Eqs. 42 and 43 is a linear function of the synaptic activity of those neurons that are presynaptic to the $i$th neuron ($\eta_i$). In

**Appendix Table 3. Parameters for the spiking network model of inspiratory rhythmogenesis.**

| Symbol | Definition | Value | Units |
|---|---|---|---|
| $N$ | network size (number of neurons) | 400 | - |
| $p$ | probability that neuron $i$ excites neuron $j$ ($i \neq j$) | 0.065 | - |
| $\delta$ | strength of recurrent excitation | 0.09 | - |
| $\bar{\eta}$ | mean excitatory drive (negative for intrinsic excitability) | $-0.0019$ | - |
| $\nu$ | variance of stochastic excitatory drive | 0.0049 | - |
| $\alpha_s$ | growth fraction for $s$ | 1 | $\mathrm{ms}^{-1}$ |
| $\tau_s$ | time constant for decay of $s$ | 10 | ms |
| $\alpha_m$ | growth fraction for $m$ | 0.4 | $\mathrm{ms}^{-1}$ |
| $\tau_m$ | time constant for decay of $m$ | 300 | ms |
| $\alpha_n$ | growth fraction for $n$ | 0.011 | $\mathrm{ms}^{-1}$ |
| $\tau_n$ | time constant for decay of $n$ | 1300 | ms |
| $\jmath_0$ | constant $\mathrm{Ca}^{2+}$ influx rate | 0.05 | $\mu\mathrm{M\,s}^{-1}$ |
| $\jmath_\eta$ | $\mathrm{Ca}^{2+}$ influx rate proportionality constant | 5 | $\mu\mathrm{M\,s}^{-1}$ |
| $\alpha_c$ | event-triggered $\mathrm{Ca}^{2+}$ influx | 0.01 | nM |
| $v_{out}$ | maximum $\mathrm{Ca}^{2+}$ efflux rate | 1 | $\mu\mathrm{M\,s}^{-1}$ |
| $\lambda_c$ | maximum $\mathrm{Ca}^{2+}$-dependent increase in synaptic drive | 0.01 | - |
| $\gamma_c$ | threshold for $\mathrm{Ca}^{2+}$-dependent increase in synaptic drive | 0.2 | $\mu$M |
| $k_c$ | reciprocal slope of $\mathrm{Ca}^{2+}$-dependent increase in synaptic drive | 0.02 | $\mu$M |

addition, the cytosolic $[\mathrm{Ca}^{2+}]$ of the $i$th neuron may be incremented when the $i$th neuron spikes,

$$\vartheta_i(t) = \pi \implies \begin{cases} c_i(t^+) = c_i(t^-) + \alpha_c \\ c_i^{tot}(t^+) = c_i^{tot}(t^-) + \alpha_c \end{cases} . \qquad [44]$$

The excitatory drive for the $i$th neuron depends on cytosolic $[\mathrm{Ca}^{2+}]$ as follows (cf. Eqs. 32 and 37),

$$\eta_i = \bar{\eta} + (\delta/\kappa_i^{in}) \sum_j a_{ij} s_j (1 - n_j) + \tilde{\eta}_i(t) + \frac{\lambda_c}{1 + e^{4(\gamma_c - c_i)/k_c}} . \qquad [45]$$

Fig. S4 shows network activity and dynamics of intracellular $[\mathrm{Ca}^{2+}]$ for the spiking model of eupnea and sigh rhythmogenesis (Eqs. 36–45) with parameters as in Appendix Tables 2 and 3. The initial values of $c_{tot}$ were normally distributed with mean 1 $\mu$M and standard deviation 0.01 $\mu$M to illustrate that synaptic activity-dependent calcium influx ($j_i^{in}$ in Eq. 43) can maintain synchronous $\mathrm{Ca}^{2+}$ oscillations. Fig. S4 may be compared to Appendix Fig. 12, which uses identical parameters save $\jmath_\eta = 0$. Comparing panel B of each figure (in particular, the third $\mathrm{Ca}^{2+}$ release event) shows how synaptic activity-dependent $\mathrm{Ca}^{2+}$ influx ($\jmath_\eta > 0$) promotes $\mathrm{Ca}^{2+}$ subsystem synchronization among the 400 neuron population.

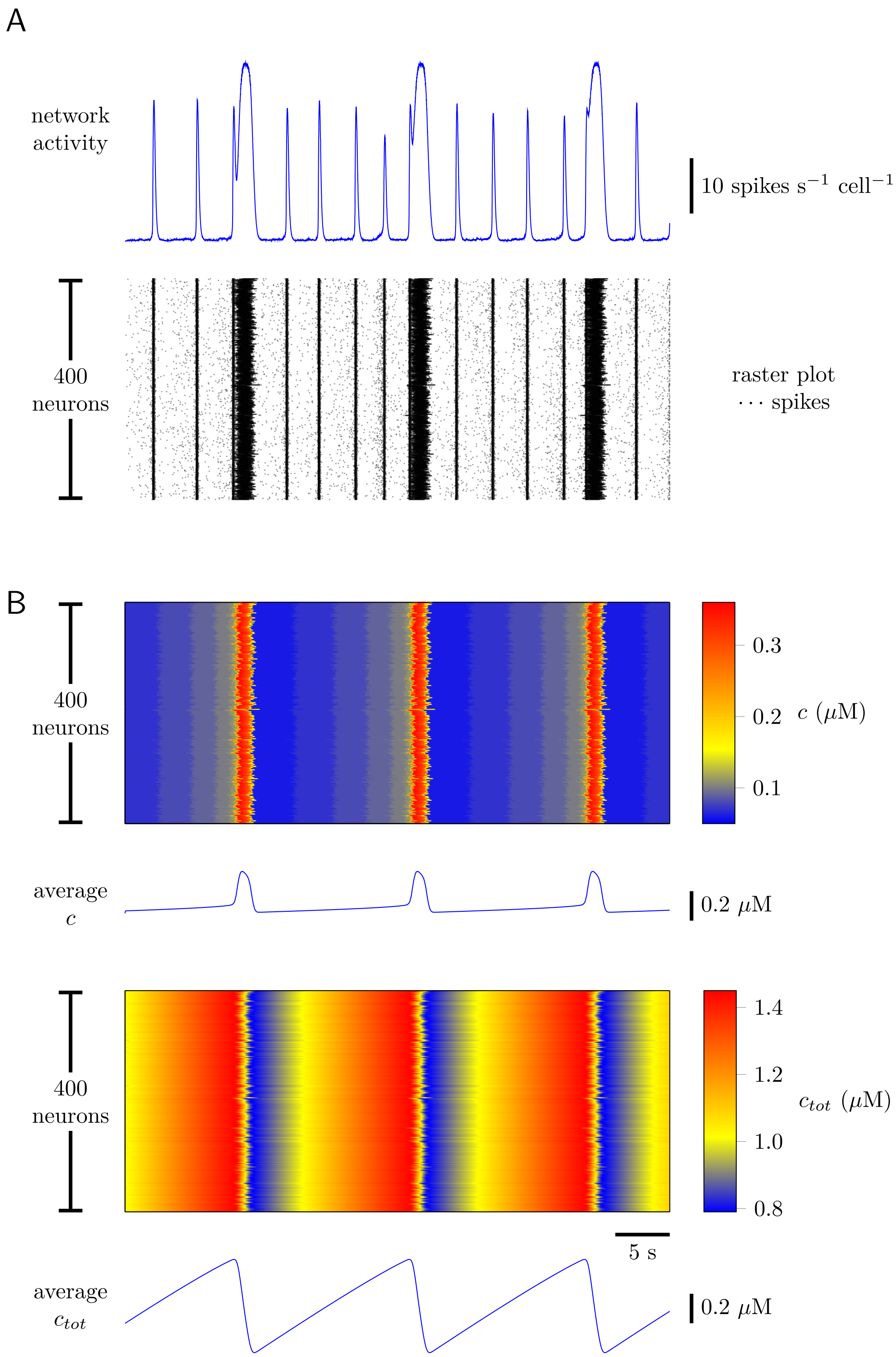


**Appendix Figure 12.** Network activity and dynamics of intracellular [Ca$^{2+}$] for a spiking model of eupnea and sigh rhythmogenesis. The network structure is Erdös-Rényi-type with 400 neurons and 6.5% probability that any given neuron is postsynaptic to any other. (A) Network activity and raster plot. (B) In the absence of synaptic activity-dependent Ca$^{2+}$ influx ($\jmath_\eta = 0$), cytosolic [Ca$^{2+}$] ($c$) and total [Ca$^{2+}$] ($c_{tot}$) remain slightly desynchronized across the 400 neurons (cf. Fig. S4). Other parameters as in top block of Appendix Table 2 and top and bottom blocks of Appendix Table 3.
